# A functional genetic landscape of antibiotic sensitivity across the pneumococcal pangenome reveals conserved and lineage-specific vulnerabilities

**DOI:** 10.64898/2026.02.26.708248

**Authors:** Bevika Sewgoolam, Axel B. Janssen, Louise S. Martin, Monica Rengifo-Gonzalez, Vincent de Bakker, Babet Rozendal, Amelieke J.H. Cremers, Jan-Willem Veening

## Abstract

The large pangenome of *Streptococcus pneumoniae* enables this opportunistic pathogen to adapt and evade antibiotic treatment. Effective treatment of pneumococcal infections requires a better understanding of the genes that modulate susceptibility to antibiotics across the pangenome. Using CRISPRi-seq, we identified genes that contribute to antibiotic sensitivity against a panel of clinically relevant antibiotics across nine pneumococcal strains with diverse resistance profiles, serotypes, and lineages. The here-generated chemical-genetics atlas revealed distinct genome-wide signatures of antibiotic stress that were specific to the antibiotic mode of action and showed both strain-specific and conserved signatures. This allowed us to identify conserved genes involved in antibiotic vulnerability and assign functions to previously uncharacterized genes. For instance, deletion of *mutS2*, which may act as a ribosome collision sensor and *spv_1295*, a conserved gene of unknown function, resulted in increased sensitivity to the macrolide azithromycin across strains, including a macrolide resistant strain, and could be potential targets for global sensitizing therapies. This work establishes a pangenome-wide framework for understanding antibiotic stress responses in *S. pneumoniae*, providing a foundation for the rational development of therapies that exploit conserved and strain-specific vulnerabilities.

## Introduction

*Streptococcus pneumoniae* (the pneumococcus) is an opportunistic pathogen that asymptomatically colonizes the human nasopharynx. However, it is also responsible for a wide range of diseases such as pneumonia, meningitis, and sepsis^1^. Despite the availability of vaccines and antibiotics, pneumococcal infections remain a leading cause of morbidity and mortality worldwide. Annually, pneumococcal infections lead to the loss of over 38 million disability-adjusted life years, and more than one million deaths, mostly in infants and the elderly^2,3^. A major challenge in controlling pneumococcal infections is the widespread emergence of antibiotic resistance^4^. Country-specific reports on isolates with reduced susceptibility to penicillin range from 14-80%^5^, with an overall global rate reported at 17.8% by the World Health Organization (WHO)^6^. Similarly, macrolide resistance has been reported to vary between 23-90% of isolates^5^, with a rate of 17.8% within the European Economic Area^7^. Fluoroquinolones however, remained effective, with resistance rates between 1-3%^5^. These high rates of antibiotic resistance are estimated to contribute to 60% of deaths caused by *S. pneumoniae*^8^. In response to the high prevalence of antibiotic resistance, the WHO has included macrolide-resistant (and previously penicillin-resistant) *S. pneumoniae* on its Bacterial Priority Pathogen List, calling for the development of new antibiotics against this species^4^.

The high antibiotic resistance rates in disease-causing *S. pneumoniae* isolates can be attributed to several factors, including the spread of high-risk (antibiotic-resistant) clones^9^, but also its high genetic adaptability. Driven by its ability to develop natural competence and transform exogenous DNA into its own genome^10,11^, *S. pneumoniae* can partake in horizontal gene transfer between itself or with other closely related species from the mitis group (e.g. *Streptococcus pseudopneumoniae* and *Streptococcus mitis)*^12,13^. This capacity facilitates the exchange of diverse traits, including capsular serotypes, virulence factors, and antibiotic resistance determinants. As a result, the pneumococcal genomic landscape is highly diversified, leading to the emergence of lineages with specialized genetic compositions that can be advantageous under adverse conditions, such as antibiotic stress^12,14,15^. These characteristics have led to a pneumococcal pangenome with a high degree of openness, in which newly sequenced genomes will continuously add new genes to the total pool^12,15–17^. In total, the size of the pneumococcal pangenome has been estimated at between 5000 and 7000 genes^12,14,17,18^. However, at the individual strain-level, an average pneumococcal genome will carry ∼2100 genes, of which ∼1350-1650 are universally present in the core genome, with ∼450-750 genes for the accessory genome^12,14,17^.

To study how the genetic variation within the pangenome influences bacterial fitness under antibiotic-induced stress, a genome-wide approach is needed. Several studies have shown that the antibacterial effects of antibiotics extend beyond the direct target of the drug and can broadly impact bacterial physiology and metabolism^19–22^. Similarly, the development of antibiotic resistance (or tolerance) can be mediated by genes outside of canonical resistance pathways. Previous studies investigating how the pneumococcal pangenome influences bacterial fitness under antibiotic-induced stress used transcriptomics and transposon-based methods such as RNA-seq and Tn-seq^20,21^. Interestingly, both these methods showed no conserved antibiotic stress response in two different pneumococcal strains to four different antibiotics^20,21^. Recent work from our lab also showed a poor correlation between gene expression and gene essentiality under infection-mimicking conditions^23^, indicating that determining antibiotic-fitness maps is more informative in predicting conserved antibiotic vulnerabilities compared to gene expression analyses. A previous study used Tn-seq to conduct a genome-wide fitness comparison across two pneumococcal strains to examine how daptomycin susceptibility in *S. pneumoniae* changes in response to gene knockouts^20^. This work showed that susceptibility changes are strain-dependent, suggesting that differences in the genomic content result in strains responding differently to the same antibiotic^20^. However, as transposon insertions result in permanent gene disruption, these studies fail to explore the role of essential genes during antibiotic treatment, as the knockout mutants of these genes will not be present in the libraries. In addition, these libraries are inherently biased towards conditions under which they were created, including the antibiotic used for selection^24^.

To address these limitations, we used CRISPR interference coupled with high-throughput sequencing (CRISPRi-seq) to study genome-wide antibiotic stress signatures. CRISPRi relies on the binding of catalytically inactive dCas9 to a sgRNA-specified chromosomal locus for transcriptional repression of the target gene^25,26^. As the expression of dCas9 is inducible, CRISPRi can be used to study the relative importance of both essential and non-essential genes, as the loss of target gene expression is not permanent. By making a pool of sgRNA clones targeting every gene in a genome, CRISPRi-seq allows for the high-throughput screening of genome-wide libraries, under diverse conditions and stresses, without prior biases^27^.

Here, we apply CRISPRi-seq to *S. pneumoniae* in a series of chemical genetic experiments under exposure of antibiotics with different mechanisms of actions^28,29^. Similar studies have shown the genome-wide genetic landscape of antibiotic sensitivity in bacteria such as *Staphylococcus aureus* and *Mycobacterium tuberculosis*^22,30,31^. However, the scope of these studies was limited, testing a small number of strains. We broadened this approach by creating CRISPRi-seq libraries in a diverse panel of nine pneumococcal strains representing the species’ wider phylogeny. Using these libraries, we identified the *S. pneumoniae* core and strain-specific essentialome. In addition, we determined how conserved and strain-specific genetic determinants modulate susceptibility to antibiotics with diverse mechanisms of action across the wider phylogeny. By generating this CRISPRi chemical-genetics atlas, we observe that the stress signatures are mode-of-action dependent, and are conserved across the pneumococcal species, contrary to previously published work^20,21^. In addition, we observe that essential genes are primarily conserved to the core genome, with only a small portion of a strain’s essential genes located in the accessory genome. Finally, we verify that several biologically relevant core genes influence antibiotic susceptibility across multiple strains using experimental approaches. The entire antibiotic-gene interaction atlas is integrated in PneumoBrowse 2, available at https://PneumoBrowse2.VeeningLab.com/^28^.

## Results

### CRISPRi-seq to determine antibiotic stress signatures in *S. pneumoniae* D39V

To identify genes that influence antibiotic-induced stress in *S. pneumoniae* D39V, we performed chemical-genetic screens by exposing a genome-wide CRISPRi library to a diverse panel of antibiotics^32,33^. This library contains a pool of mutants harboring 1498 unique sgRNAs that target 98% of the annotated genetic elements under the control of a constitutive promoter, and *dcas9* under the control of an isopropyl β-D-1-thiogalactopyranoside (IPTG) inducible promoter. Each operon is targeted by one sgRNA (“D39V operon-level library”), providing us with a genome-wide view of operons contributing to increased or decreased drug-sensitivity. For these experiments, a broad panel of 15 antibiotics, across five different classes and four different modes of actions, was selected: cephalosporins, fluoroquinolones, macrolides, penicillins, and rifamycins (Figure 1A, Supplementary Data 1). Of these, the β-lactam antibiotics (i.e., cephalosporins and penicillins) share a similar mechanism of action by inhibiting penicillin-binding proteins, although they differ in their affinity for these cell-wall synthesizing enzymes^34^. Both clinically relevant antibiotics currently used to treat pneumococcal infections and antibiotics not typically used in treatment were included in this study. The β-lactam antibiotics (and amoxicillin in particular) are clinically relevant for pneumococcal infections^35^. Antibiotics such as the rifamycins are not generally used in the treatment of pneumococcal infections, but they are used in the treatment of pulmonary diseases such as tuberculosis, where secondary pneumococcal infections can occur, potentially exposing pneumococci to these drugs^36^. The inclusion of such antibiotic classes may not be directly relevant for pneumococcal infection-treatments but provides insights into the unique antibiotic-induced stress signatures of different classes of antibiotics with varying modes of action.

**Figure 1.**
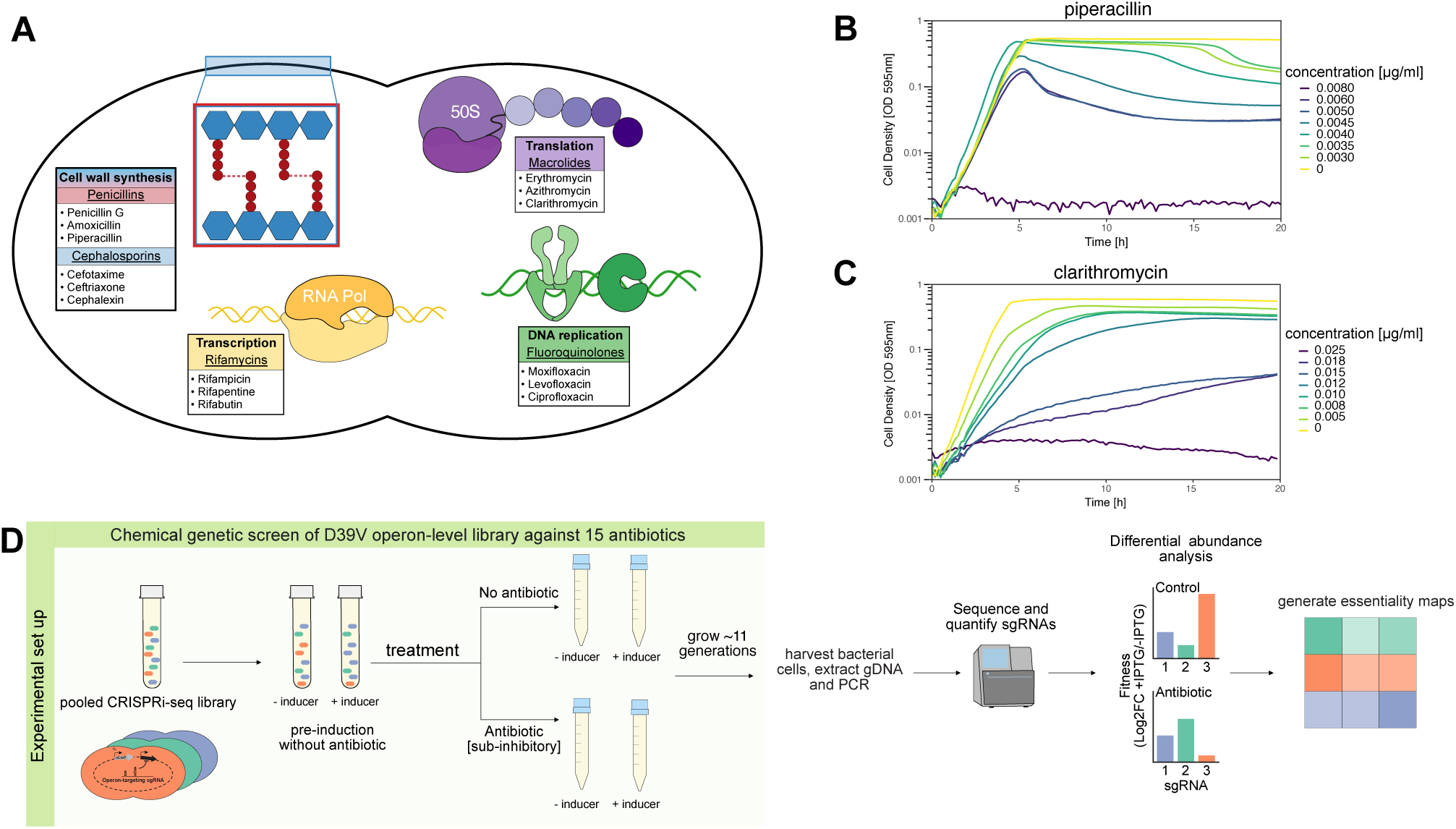
Experimental setup of CRISPRi-seq chemical genetic screens in *S. pneumoniae* D39V. **A)** Overview and schematic representation of the mode of action and targeted processes of the five antibiotic classes used in the screen. Representative growth curves used to determine sub-inhibitory concentrations are shown for **B)** piperacillin and **C)** clarithromycin. Final concentrations selected from the growth curves were 0.004 and 0.012 μg/ml, respectively (Supplementary Data 1). **D)** Schematic representation of the experimental workflow for the chemical genetic screens using the operon-level CRISPRi-seq library in *S. pneumoniae* D39V against a panel of 15 different antibiotics.

For these screens, we required a suitable sub-inhibitory antibiotic concentration to exert sufficient antibiotic pressure and substantially impair bacterial growth, without imposing a bottleneck that would be too strict as to hinder downstream differential abundance analyses. To determine the sub-inhibitory concentrations to be used in the CRISPRi-seq screens, we used growth curves of *S. pneumoniae* D39V exposed to varying antibiotic concentrations, spanning minimum to maximum growth inhibition. Representative growth curves from piperacillin (Figure 1B, 0.004 μg/ml) and clarithromycin (Figure 1C, 0.012 μg/ml) show the impaired growth at these concentrations, compared to antibiotic-free control conditions. After selection of the sub-inhibitory antibiotic concentrations (Supplementary Data 1), the screens were conducted by culturing the CRISPRi-seq library in one of four conditions: I) minus IPTG, minus antibiotic; II) plus IPTG, minus antibiotic; III) minus IPTG, plus antibiotic; or IV) plus

IPTG, plus antibiotic. After culturing, cells were harvested, gDNA was extracted, and sgRNAs were amplified and sequenced (Figure 1D).

### Antibiotic stress signatures are dependent on antibiotic mechanism of action

After sequencing, quantification, and normalization of sgRNA abundances (Supplementary Data 2), we explored how the overall composition of sgRNAs changed in response to the sub-inhibitory antibiotic concentrations. Principal component analysis (PCA) of the sgRNA abundances showed that all uninduced samples (-IPTG) were similar in their overall sgRNA composition, but distinct from induced (+IPTG) samples (Figure 2A). The similarity of the uninduced samples confirmed that the concentrations used did not impose a significant bottleneck on the library, and the tight control of dCas9 expression. The antibiotic-exposed, IPTG-induced samples formed distinct, class-dependent clusters, reflecting mechanism of action-dependent antibiotic stress signatures (Figure 2A).

**Figure 2.**
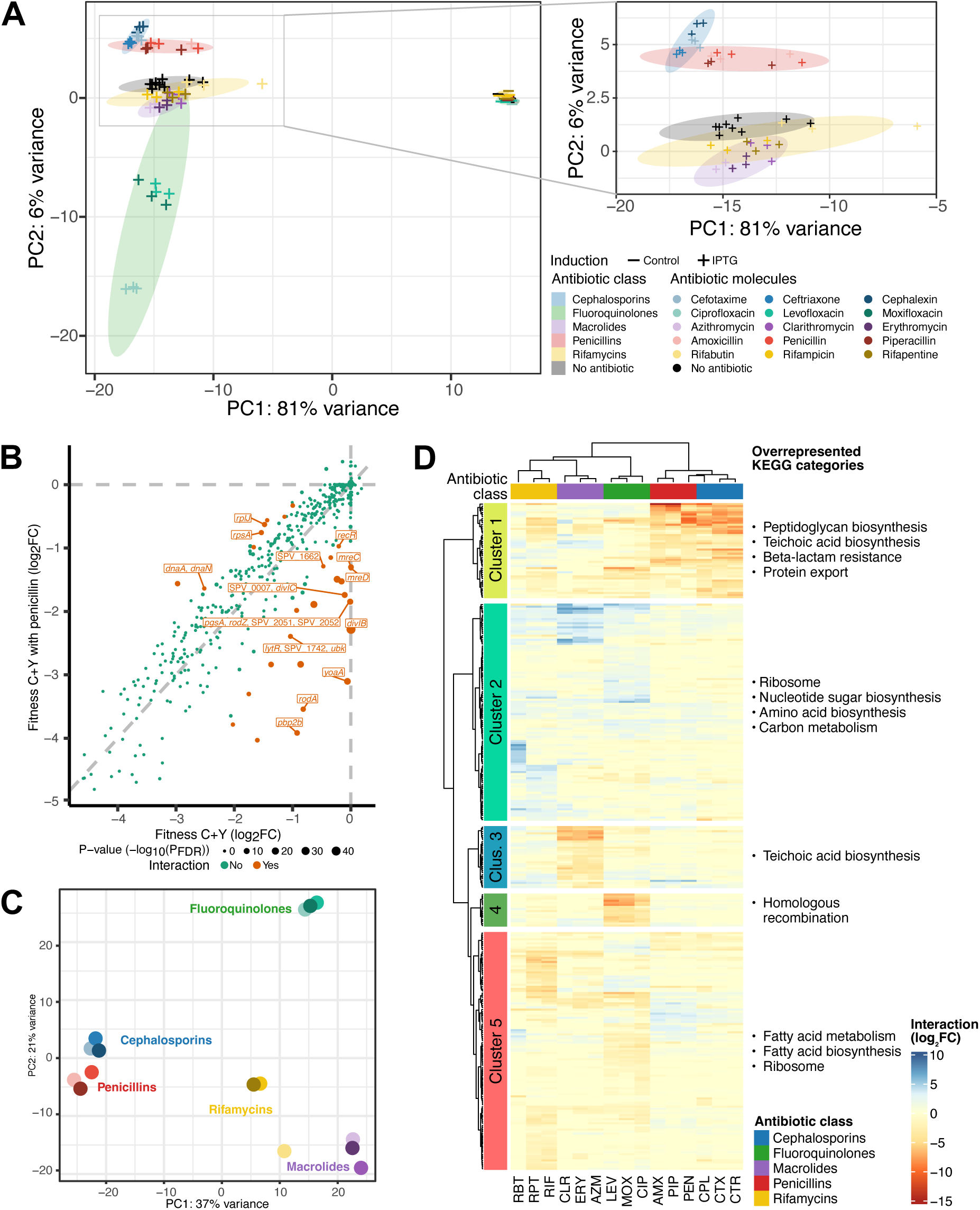
Genome-wide antibiotic stress signatures of *S. pneumoniae* D39V reflect antibiotic mode of action. **A)** PCA of overall sgRNA composition shows that 81% of variance is explained by the *dcas9*-induction status. Antibiotic exposure explains 6% of the variance in the data. Ellipses represent 95%-confidence intervals based on class-level analyses (nine datapoints for each class, nine for control conditions). A magnified view of the main plot depicts the clustering of antibiotic classes. **B)** Scatter plot of scaled log2-transformed fold changes (log2FCs) showing the effect of antibiotic addition (penicillin shown here) to C+Y medium on fitness upon knockdown. Interaction is determined by cutoff values for interaction log2FCs (absolute value > 1), and the associated adjusted P-value (< 0.05). All essentiality data is available in Supplementary Data 4. **C)** PCA plot of the scaled interaction log2FCs. Like the overall sgRNA composition, the interaction results show specific clustering based on the mode of action of the antibiotic class. The legend from panel A applies here as well. **D)** Heatmap depicting scaled interaction fitness scores of sgRNA targets with significant differential fitness upon treatment within at least one antibiotic (n=312). Like the PCA plot in panel E, the column’s hierarchical clustering (Ward’s method, implemented through the “ward.D” method in R’s hclust function, using Euclidean distance), shows similarity based on antibiotic mode of action. Row clustering (same methods), shows clustering into distinct clusters. The overrepresented KEGG pathways per cluster are indicated. Antibiotic abbreviations: AMX, amoxicillin; AZM, azithromycin; CIP, ciprofloxacin; CLR, clarithromycin; CPL, cephalexin; CTX, cefotaxime; CTR, ceftriaxone; ERY, erythromycin; LEV, levofloxacin; MOX, moxifloxacin; PIP, piperacillin; RIF, rifampicin; RBT, rifabutin; RPT, rifapentine.

After the initial analysis of sgRNA composition, we compared the sgRNA counts from each antibiotic to control conditions in a differential fitness analysis. This comparison reveals each operon’s differential fitness upon knockdown across the 15 antibiotics. Of note, direct targets of the antibiotics (e.g., *rpoB* for rifamycin) do not necessarily appear in the differential fitness analysis as hits, as these targets were baseline essential without addition of antibiotics (Supplementary Data 3). Treatment with antibiotics results in enrichment or depletion of specific sgRNAs, indicating that these targets contributed to increased or decreased fitness respectively, in the presence of antibiotic. For example, within the penicillin dataset, the knockdowns of *pbp2b*, *rodA*, and *divIB* were more detrimental for survival under sub-inhibitory concentrations of the antibiotic, as they had a greater negative log_2_FC in presence of the antibiotic, than under control conditions (Figure 2B). These genes all serve a role in cell wall synthesis and thus reflect the mode of action of penicillin (i.e., inhibition of the cross-linking of the cell wall peptidoglycans by binding to penicillin-binding proteins).

Similarly, *S. pneumoniae* D39V treated with cephalexin identified increased vulnerabilities in genes associated with cell morphology and division: *pbp1a*, *pbp2b,* and *murA1* involved in peptidoglycan synthesis; *rodA*, *rodZ*, *mreC,* and *mreD* involved in establishing cell shape, and *divIB, and divIC* involved in cell division (Supplementary Figure 1, Supplementary Data 4). In addition, the most prominent hits for ciprofloxacin were genes involved in DNA repair and recombination, as described previously^37^. When the interaction values of only those sgRNAs that had a significant interaction (absolute log_2_FC > 1, P-value < 0.05) in presence of one or more antibiotics were considered and summarized using a PCA plot, the mode of action-dependent clustering is evident (Figure 2C). These differential fitness analyses thus identify genes with functions specific to the mode of action of each antibiotic class (Supplementary Figure 1, Supplementary Data 4).

The observed pattern of mode of action-dependent antibiotic stress signatures was further visualized using clustered heatmaps of those sgRNAs that had a significant interaction in presence of one or more antibiotic (Figure 2D). For each operon, the relative gain (blue), or loss (red) of fitness upon addition of an antibiotic was visualized, revealing associations between antibiotics and genetic elements. We observed clear clustering of antibiotics based on their mechanism of action, similar to the PCA analysis. In addition, this analysis allowed us to visually identify genes driving the clustering. For the β-lactam antibiotics, the genes that conferred a fitness defect upon knockdown (Cluster 1) have been previously described to play a role in β-lactam resistance (e.g. *divIB*^38^, *murA*^39^, and *pbp2b*^40^) and play a role in cell wall-related processes, according to a KEGG pathway analysis. Like the individual differential fitness analyses (Figure 2B), the mechanism of action of the antibiotics are thus reflected by the genes whose knockdown confers a fitness change in the presence of that antibiotic. Within Cluster 2, different ribosomal, nucleotide sugar / amino acid biosynthesis, and carbon metabolism genes cluster together. The knockdown of these genes is observed to be relatively beneficial in the presence of fluoroquinolones, macrolides and rifamycins. One explanation for this could be that knockdown of these genes leads to overall reduced growth, which may curb the antibiotics’ effects and may implicate these genes in the generation of antibiotic persistence^41^. The knockdown of teichoic acid biosynthesis genes (Cluster 3) results in a fitness defect when exposed to macrolides, whilst genes involved in homologous recombination are important for survival in conditions with fluoroquinolones (Cluster 4). The genes represented in Cluster 5 do not seem to be particularly important or detrimental for a specific condition, although fatty acid-pathways and ribosome-related genes are overrepresented.

Full results of the chemical genetic screens using the D39V operon-level library can be found in Supplementary Data 2 (raw and normalized sgRNA counts), Supplementary Data 3 (essentiality results), and Supplementary Data 4 (interaction results).

### Core genome essentiality over multiple timepoints

The clear antibiotic mode of action-dependent stress signatures observed in *S. pneumoniae* strain D39V prompted us to further investigate whether a similar antibiotic stress signature is present across the wider *S. pneumoniae* species. For this, we created tailored sgRNA libraries in nine phylogenetically diverse strains, including well-studied strains such D39V and TIGR4^42,43^, strains that are known for their proficient ability to colonize the nasopharynx (EF3030 and BHN418)^44,45^, and strains that were isolated from patients in recent studies (PBCN0272, PBCN0364)^46^ (Figure 3A, Table 1). These strains covered diverse characteristics in the pneumococcal species, and their core genome (1606 protein-encoding genes) was of similar size compared to the 38 strains that represented the full pneumococcal phylogeny (1535 genes, Figure 3B)^47^. In each strain, we cloned *dcas9* under the anhydrotetracycline (aTc) and doxycycline-inducible P*tet* promoter at an ectopic locus (downstream of the conserved *bgaA* gene) and confirmed CRISPRi functionality by cloning the sgRNAs to target *axe1* and *tarI* (Supplementary Figure 2). Next, these strains were used to create strain-specific genome-wide sgRNA libraries to enable CRISPRi-seq. However, as an operon-level design, as used in the D39V library before, is difficult to establish solely based on genomic sequence analysis, we instead targeted every gene individually (“gene-level libraries)”. Using a recently developed computational method^48^, we designed sgRNAs (2056-2230 per strain) to target every predicted coding feature in the genomes of these strains (Supplementary Data 5). These sgRNAs were subsequently synthesized as a pooled oligonucleotide library, from which strain-specific portions were amplified for downstream use. Through a high-throughput cloning method using Esp3I-mediated Golden Gate cloning, we succeeded in cloning all strain-specific sgRNAs, whilst limiting the portion of cells encoding foreign sgRNA sequences to 0.036-0.054% of the libraries (Supplementary Figure 3).

**Figure 3.**
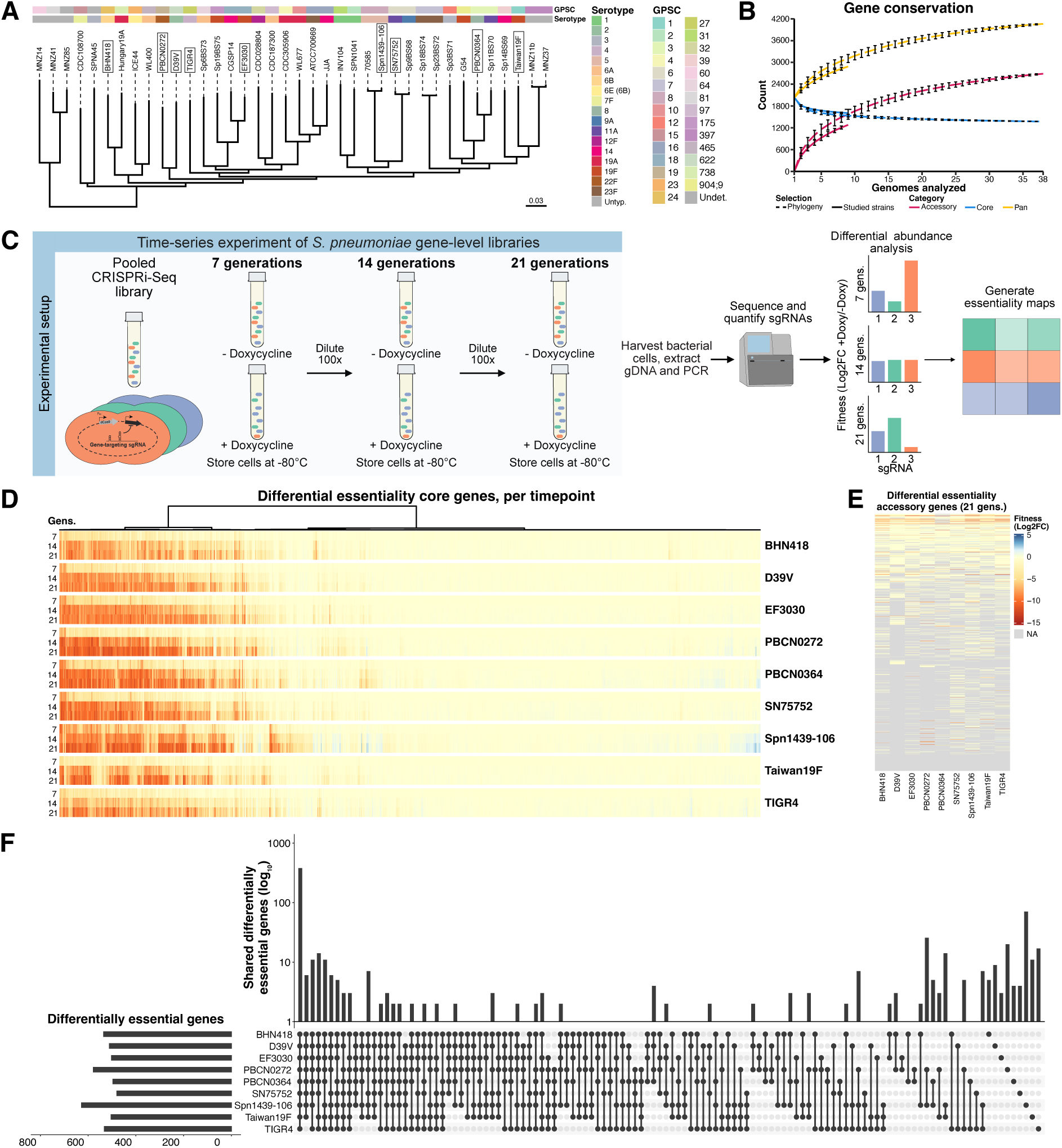
Multi-strain CRISPRi-seq reveals gene essentiality across the pneumococcal phylogeny. **A)** Phylogenetic tree indicating phylogenetic standings of the strains used in this work. Besides the nine genome of the strains used in this work, the genomes of 29 reference strains were included in the analysis, to provide for the overall pneumococcal phylogenetic structure^47,99^. The genomes of the phylogenetic distinct naturally nonencapsulated *S. pneumoniae* strains MNZ14, MNZ41, and MNZ85 perform as outgroup. This SNP-based tree is based on a recombination-corrected core genome alignment measuring 1.61 - 1.84 Mbp to the reference strain (D39V). For each strain, the capsular serotype and Global Pneumococcal Sequence Cluster (GPSC) as determined by Pathogenwatch are indicated^100^. Untypeable (untyp.), and undetermined (undet.) serotypes and GPSCs are greyed out. The phylogenetic distance is based on a 73.4 kbp polymorphic site alignment. The genomes of strains used in this work are highlighted within a box. **B)** Analysis of the genetic content shows the similarity in core genome size in between the full phylogeny pictured in panel A, and the studied strains. At each number of genomes analyzed, a combination of that number of genomes was subset from the available genomes. For the studied strains, all available combinations were analyzed, whilst for the full phylogeny, a maximum of 1000 unique, random combinations were analyzed. At nine analyzed genomes, the studied strains have a core genome of 1606, whilst the full phylogeny had a core genome of 1535 protein-encoding genes. In this analysis, only protein-encoding genes are considered by Panaroo^83^. **C)** Schematic representation of the experimental workflow for the time-series screen using the gene-level CRISPRi-seq libraries in the phylogenetically distinct *S. pneumoniae* over multiple generation timepoints. **D)** Differential fitness analysis of 1549 core genes show the conserved gene necessity in the core genome, across timepoints. At 7, 14, and 21 generations after start of dCas9-induction, respectively 170, 334, and 376 sgRNAs were determined to be universally differentially present at those timepoints. **E)** In contrast to the core genome, the accessory genome shows less differentially present sgRNAs. After 21 generations, an average of 43 accessory genes were putatively essential per genome, on an average of 431 accessory genes. Genes absent in a specified genome are greyed out. **F)** Overlap in differentially essential core genes between genomes after 21 generations of dCas9-induction. After 21 generations, 376 genes were determined to be universally differentially present. Outside of these universal genes, a total of 402 genes showed reduced fitness upon depletion in one or more strains, with 71 genes differentially essential in Spn1439-106.

**Table 1.** Strains used in this study.

Strains used in this study
| Strain | Serotype | Global Pneumococcal Sequence Cluster | Significance | Accession number | Reference | Remark |
| --- | --- | --- | --- | --- | --- | --- |
| **BHN418** | 6B | 24 | Nasopharyngeal carriage isolate | GCF_000495375.1 | 44 |  |
| **D39V** | 2 | 622 | Model strain | CP027540.1 | 42 |  |
| **EF3030** | 19F | 18 | Nasopharyngeal carriage isolate | NZ_CP035897.1 | 45 |  |
| **PBCN0272** | 22F | 19 | Blood culture isolate | ERS225756 | 46 |  |
| **PBCN0364** | 8 | 3 | Blood culture isolate | ERS225822 | 46 |  |
| **SN75752** | 11A | 6 | Blood culture isolate.<br>Amoxicillin resistant. | CP089949.1 | 40 | Named "11A" in reference |
| **Spn1439-106** | 19A | 8 | PMEN19 reference strain. | CP155533.1 | 101 |  |
| **TIGR4** | 4 | 27 | Model strain | NC_003028.3 | 43 |  |
| **Taiwan19F** | 19F | 1 | PMEN14 reference strain.<br>Penicillin and macrolide resistant. | NC_012469.1 | 102 | Named "TW31" in PneumoBrowse 2 |

To assess the quality of these libraries, we performed a time-series experiment. By collecting cells after 7, 14, and 21 generations of exponential growth in the presence (or absence) of 20 ng/ml of doxycycline to induce dCas9 (Figure 3C), we assessed the genome-wide gene essentiality across the phylogeny under standard laboratory culture conditions (C+Y medium, 37°C). By assessing the differential fitness at different timepoints, we aimed to determine whether essential processes within *S. pneumoniae* could be ranked according to the fitness defect that transcriptional inhibition of involved genes would yield^49–51^. Our phylogeny-wide analysis showed that the gene necessity in these nine strains is conserved, both at different timepoints, and across the assessed core genome (n=1549) (Figure 3D; Supplementary Figure 4). After 7 generations, 170 sgRNAs were determined to have a differential abundance, whilst after 14 generations 334 sgRNAs were altered. After 21 generations 376 sgRNAs had a differential abundance, or 23.4% of the sgRNAs in the core genome. Over the different timepoints, genes assigned to Clusters of Orthologous Genes (COG) functional group J (translation, ribosomal structure and biogenesis) made up the largest part of the genes that showed reduced fitness (Supplementary Figure 5).

In contrast to the core genome, only 1/10^th^ of the accessory genome of a given strain showed reduced fitness upon CRISPRi knockdown under these laboratory conditions (Figure 3E). Within the core genome, half of the putative essential genes showed reduced fitness in all strains (n=376/778), although a significant portion was specifically differentially essential (n=144), or non-essential (n=59), in a specific strain (Figure 3F, Supplementary Figure 6). The observation of strain-specific gene essentiality was confirmed by cloning of select sgRNAs and growing them individually. Strains with sgRNAs targeting *upp* and (the ortholog of) *spv_0645* showed selective essentiality in strains D39V and Spn1439-106 respectively, whilst selective non-essentiality was observed for *accB* in strain SN75752 (Supplementary Figure 2).

Full results on the raw counts, normalized counts, and essentiality on each timepoint by strain, can be found be in Supplementary Data 6, Supplementary Data 7, and Supplementary Data 8 respectively. Notably, the identified essentialomes in strains D39V, Taiwan19F, and TIGR4 correspond well with previously established essentialomes based on Tn-seq approaches (Supplementary Data 9)^18,32,52^, demonstrating the high quality of the here-established CRISPRi libraries.

### Multi-strain chemical-genetics shows conserved species-wide antibiotic vulnerabilities dependent on antibiotic mode of action

To identify antibiotic stress signatures across the wider *S. pneumoniae* species, we exposed the nine gene-level CRISPRi libraries to antibiotics with different mechanisms of action. For these experiments, we selected four clinically relevant antibiotics of four different classes: amoxicillin (β-lactam), azithromycin (macrolide), levofloxacin (fluoroquinolone), and linezolid (oxazolidinone) (Figure 4A). The selection of a single antibiotic from each class was informed by results of the D39V operon-level CRISPRi-seq screening which indicated that gene fitness was similar between antibiotics of the same class (Figure 2D). The inclusion of linezolid (which targets protein synthesis by binding to the 23S ribosomal subunit) was motivated by its excellent pharmacokinetic and pharmacodynamic properties and its potential use for the treatment of Gram-positive bacterial infections in the future^53–56^. In addition, the inclusion of two antibiotics that have a similar mechanism of action (linezolid and azithromycin; inhibition of protein synthesis through binding of ribosomal subunits), should further confirm our previous observations that antibiotic stress signatures are dependent on the mode of action.

**Figure 4.**
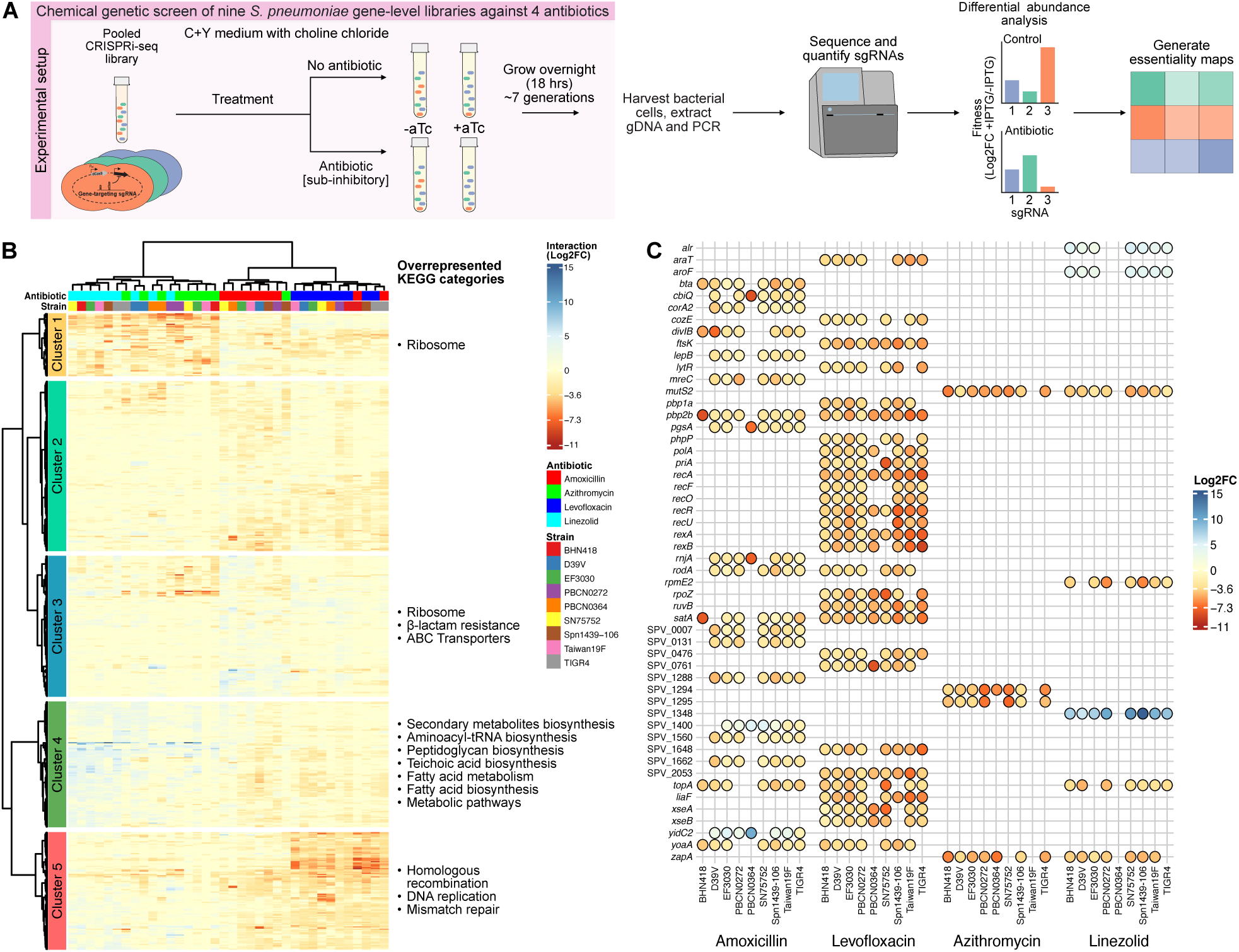
Mode of action-dependent antibiotic stress signatures are conserved across the *S. pneumoniae* core genome. **A)** Schematic representation of the overnight experimental setup for the multi-strain chemical genetic CRISPRi-seq screen. For this screen, C+Y medium was supplemented with choline chloride (10 mg/ml) to prevent LytA-mediated autolysis during overnight growth. **B)** Heatmap of scaled log2FC scores of sgRNA targets with significant differential fitness upon treatment in at least three datasets (combination of strain and antibiotic). Genes that resulted in fitness loss upon knockdown are represented in red and genes that resulted in a fitness gain upon knockdown are represented in blue. Strain name and antibiotic treatment are represented by specified colors. Only genes that have a homologue in all 9 strains were used in this analysis. Hierarchical clustering was performed with Ward’s method, implemented through the “ward.D” method in R’s hclust function, using Euclidean distance. Overrepresented KEGG categories per cluster were determined by analyzing the genes represented in that cluster for overrepresented KEGG clusters using clusterProfiler^97^. No specific KEGG clusters were returned for Cluster 2. **C)** Comparative bubble plot of gene fitness across four antibiotics and nine strains. Only conserved genes that showed a statistically significant differential fitness effect in at least seven strains upon antibiotic treatment are considered in this comparison.

We first explored the essentiality within each of the nine strains treated with sub-inhibitory concentrations of one of the four antibiotics (Supplementary Figure 7 and Supplementary Figure 8, Supplementary Data 10). We then compared the differential fitness across the core genome of the nine strains and four antibiotics (Supplementary Data 11). This showed that antibiotic stress signatures and vulnerabilities were conserved. The mode of action-dependent clustering of the antibiotic stress signatures is evident through the close clustering of the azithromycin and linezolid-exposed samples in a single branch (Figure 4B). The amoxicillin and levofloxacin samples however, clustered in two separate branches. These results contradict previous Tn-seq mediated studies, in which no pattern of conserved gene essentiality was observed^20^. This contradiction presumably stems from the different nature of gene suppression mediation by Tn-seq compared to CRISPRi-seq. Since transposon insertion leads to a full knockout of the gene, the inactivation is permanent in nature. In contrast, CRISPRi-seq, permits inducible knockdown of specific genes. This allowed for the screening of essential genes, which are well-known to be involved in the mechanisms targeted by the antibiotics.

The observation that genes with differential fitness corresponded directly to the process that the antibiotic targeted (Figure 4C), was comparable to the results of the operon-level D39V chemical genetic screens (Figure 2D). Under influence of levofloxacin, genes involved in DNA repair and recombination became more important such as *rexAB* (involved in DNA damage response), *rec* genes (broadly involved in DNA recombination) and *priA* (DNA helicase)^32,37,57^ (Figure 4C, Supplementary Figure 9). Interestingly, in six out of nine strains, *liaF* appeared as a significant hit, resulting in a fitness defect upon knockdown in the presence of levofloxacin, consistent with previous findings (Figure 4C)^37^. The observed fitness defect is likely due to downregulation of the downstream located *liaS* causing upregulation of the LiaR regulon, which renders pneumococci more susceptible to fluoroquinolone antibiotics^37^.

The knockdown of genes involved in cell wall synthesis resulted in a fitness defect under amoxicillin stress (such as *divIB*, *mreC, pbp2b*, *rodA*, *yoaA*), while hits such as *yidC2*, *amiA* and *eloR* resulted in a fitness gain in the presence of amoxicillin across the phylogeny (Figure 4C, Supplementary Figure 9). Knockdown of *amiA* resulting in a fitness advantage in the presence of amoxicillin corroborated a previous report demonstrating that disruption of the *ami*-operon, encoding a peptide transporter, results in reduced antibiotic sensitivity^19^. The knockdown of *eloR* (*khpA*), an RNA-binding protein and regulator of cell elongation and division in *S. pneumoniae*, resulted in a fitness advantage in the presence of amoxicillin. This observation is consistent with previous studies that found the deletion of *eloR* rendered *pbp2b* and *rodA* redundant, with cells displaying a reduced growth rate^58–60^. (Supplementary Figure 9). Since β-lactams like amoxicillin target actively growing and dividing cells, these slower-growing cells are possibly less sensitive to the antibiotic.

Interestingly, amoxicillin-resistant strains SN75752 and Taiwan19F show a fitness advantage upon gene knockdown of the non-essential penicillin-binding protein *pbp2a* and the serine/threonine phosphatase *phpP* (Supplementary Figure 9). In contrast, deletion of these genes resulted in a fitness disadvantage in all the other amoxicillin-susceptible strains. PhpP dephosphorylates StkP, a key regulator of cell division genes^61,62^. One hypothesis is that deleting *phpP* causes StkP hyperactivity, which compensates for fitness defects in amoxicillin-resistant strains by enhancing cell wall synthesis. However, in amoxicillin-susceptible strains, this disrupts normal division, leading to morphological defects and reduced fitness. This may suggest the presence of an amoxicillin-resistant strain-specific signature; however, this remains to be validated.

Full results on the raw counts, normalized counts, and essentiality in each antibiotic condition by strain, can be found be in Supplementary Data 12, Supplementary Data 13, and Supplementary Table 14, respectively.

### Inferred functional roles of hypothetical proteins associated with macrolide vulnerability

Several genes were identified as essential under azithromycin stress in most strains including *mutS2* (annotated as being involved in mismatch DNA repair), *zapA/B* (divisome proteins) and *spv_1295* (a putative Zn^2+^-binding, hemolysin transmembrane protein)^63^ (Figure 4C, Figures 5A, 5B, Supplementary Figure 7). We validated the CRISPRi-seq data by making *mutS2* and *spv_1295* deletions in strains D39V and PBCN0272 (Figures 5C, 5D), and while we attempted to make a *mutS2* deletion mutant in Taiwan 19F, the gene was found to be essential in this strain. As shown in Figure 5C and 5D, Δ*spv_1295* and Δ*mutS2* mutants in strains D39V and PBCN0272 treated with sub-inhibitory concentrations of azithromycin were more vulnerable, resulting in cell death compared to wild type. The Δ*spv_1295* mutant in T19F resulted in a slight growth defect in the presence of azithromycin (Figure 5E). In addition, these mutants were more vulnerable to other macrolides such as clarithromycin and erythromycin, as well as linezolid, as predicted by CRISPRi data (Supplementary Figure 9 & 10).

**Figure 5.**
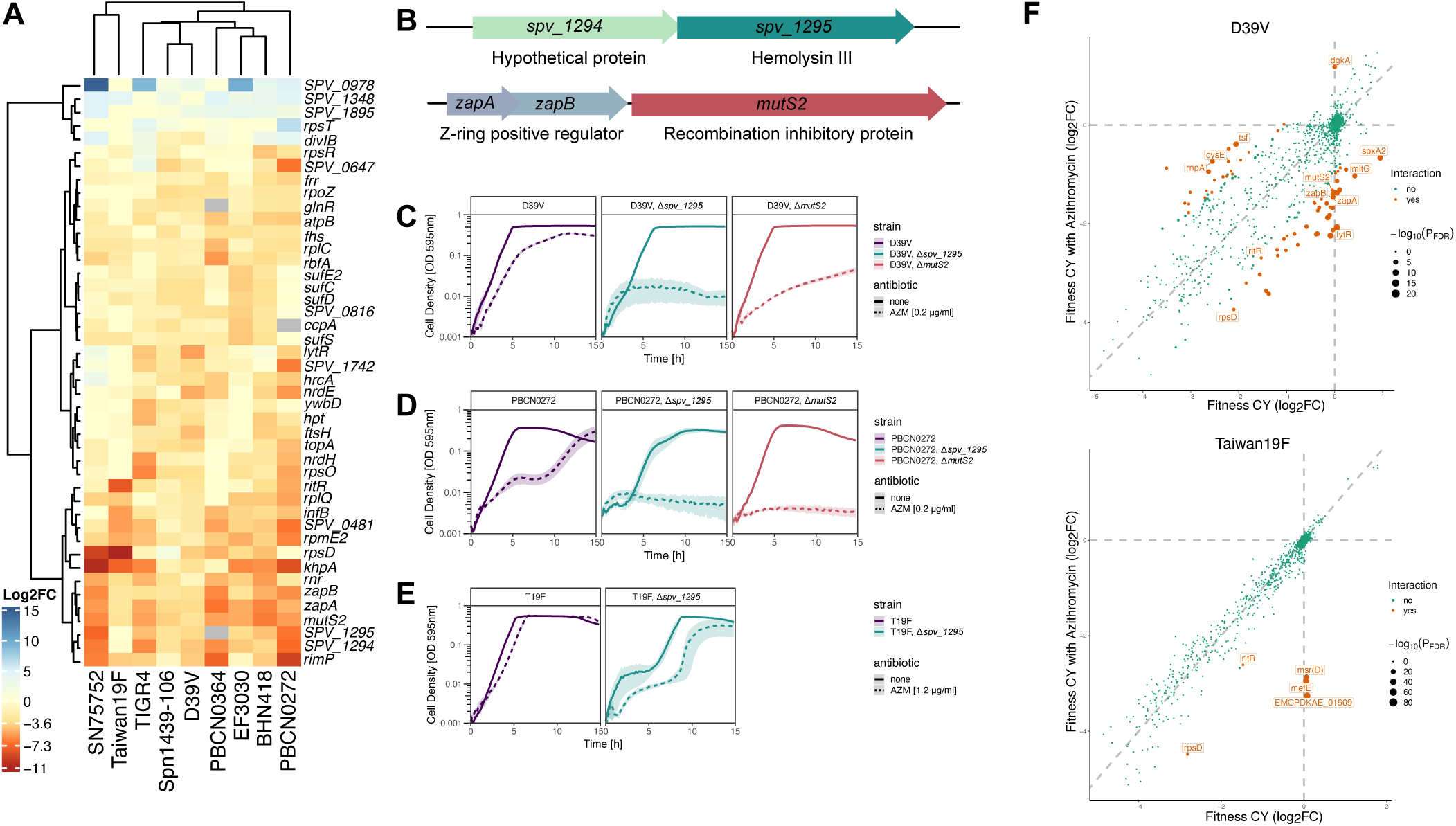
Genetic determinants of azithromycin sensitivity across pneumococcal strains. **(A)** Heatmap of all sgRNAs showing a significant differential fitness effect upon azithromycin treatment compared with antibiotic-free control in at least three out of nine strains. **B)** Schematic representation of genomic context of gene deletions for genes *spv_1295* and *mutS2*. **C)** and **D)** Growth curves of wild-type and deletion mutants (Δ*spv_1295* and Δ*mutS2*) in the macrolide-sensitive strains **(C)** D39V and **(D)** PBCN0272 treated with sub-inhibitory concentrations of azithromycin. Both deletions result in killing upon azithromycin exposure compared with wild type. **E)** Growth curves in the macrolide-resistant strain Taiwan19F (T19F), showing that deletion of *spv_1295* increases azithromycin sensitivity but does not result in cell death. Growth curve data represent the mean ± SEM of three biological replicates. **F)** Interaction analysis scatter plots for D39V (top) and Taiwan19F (bottom), showing genes with significant differential fitness effects upon CRISPRi knockdown with azithromycin relative to control samples. Fitness scores are measured by the log2FC of sgRNA counts between induced and uninduced samples. Points in orange indicate a significant interaction between induction and antibiotic treatment. Point size indicates the - log10 transformed adjusted P-value. In Taiwan19F, *msrD*, *mefE* and *EMCPDKAE_01909* are significant hits and are genes associated with macrolide resistance.

A study in *B. subtilis* showed that *mutS2*, generally known to be involved in DNA repair, has an additional role in resolving ribosome stalling and a *mutS2* deletion strain exhibited increased sensitivity to erythromycin^64^. Macrolides (azithromycin, erythromycin) and linezolid bind to the 50S ribosomal subunit, causing ribosome stalling by blocking peptide elongation. When *mutS2* is deleted, the ability to resolve ribosome collisions is potentially lost, leading to an accumulation of stalled ribosomes, increased translational stress and consequent sensitivity to macrolides^64^. Why *zapA/B* becomes particularly important during perturbations in protein synthesis remains to be tested but is likely due to their upstream location to *mutS2* (Figure 5B) and do not represent bona fide hits. Interestingly, Tn-seq screens in *S. aureus* also identified that inactivation of the genes *mutS2* and *zapA* increased susceptibility to linezolid, making these interesting targets in Gram-positive bacteria^65^.

The exact mechanism through which SPV_1295 acts, remains to be unraveled but clean knock out assays demonstrate that the absence of *spv_1295* renders D39V and PBCN0272 more susceptible to azithromycin, and also the azithromycin-resistant strain Taiwan 19F, however to a lesser degree (Figure 5E). SPV_1295 shares significant identity (>50%) with proteins homologous to PAQR (Progestin and AdipoQ Receptor) family membrane homeostasis proteins such as TrhA found in Staphylococci and Fusobacterium^28^. TrhA of *E. coli* and *B. subtilis* were shown to regulate membrane energetics homeostasis^66^. Thus, a tempting hypothesis is that downregulation of *spv_1295* in *S. pneumoniae* leads to increased antibiotic influx due to perturbed membrane permeability. Whether this hypothesis is correct and what the exact function is of SPV_1295 needs further investigation.

Interestingly, even strains that have a genetic predisposition to resist the effects of these antibiotics (e.g., Taiwan19F against azithromycin, and SN75752 against amoxicillin), clustered with the other strains (Figure 4C, Supplementary Figure 9C), indicating that the antibiotic stress signature functions independent of any specific resistance mechanism, even though the changes may not be defined as significant. This suggests that targeting factors such as MutS2 and SPV_1295 could be a promising strategy for adjunct antibiotic therapy, potentially restoring sensitivity in resistant strains. Strikingly, knockdown of *mefE* and *msrD* in strain Taiwan19F resulted in increased azithromycin susceptibility (Figure 5F). MefE and MsrD encode components of a well-characterized macrolide efflux system^67^, demonstrating that CRISPRi-seq can pick up known antibiotic-resistance determinants. The sgRNA EMCPDKAE_01909 which targets a hypothetical gene directly upstream of *mefE* also showed reduced fitness upon macrolide exposure (Figure 5F), suggesting that this is due to polar CRISPRi effects on *mefE*.

## Discussion

The current lack in development of novel antibiotic compounds calls for significant efforts to determine suitable novel targets. To address this issue, we utilized a chemical-genetic CRISPRi-seq approach across multiple pneumococcal strains to determine the genome-wide factors that affect bacterial fitness in the presence of antibiotics. Here, we provide a more comprehensive understanding of the genetic factors influencing pneumococcal survival and stress response to antibiotics needed to further guide drug development. The data generated here serves as a source to aid in future vaccine and antibiotic design. The entire antibiotic-gene interaction atlas is integrated in PneumoBrowse 2, available at https://PneumoBrowse2.VeeningLab.com/^28^.

We provide proof-of-concept in the well-characterized model strain D39V that CRISPRi-seq can be used to reveal genes that have differential essentiality during antibiotic stress. This can be further extrapolated to generate genome-wide antibiotic-specific genetic signatures which correspond to antibiotic mode of action and class. We identified numerous genes influencing antibiotic sensitivity. For example, knockdown of well-characterized genes, such as cell wall synthesis genes (*divIBC*, *yoaA*, *rodZ*, *murA1*) increased β-lactam sensitivity, while knockdown of the *ami* antibiotic importer (*ami*-operon genes) decreased sensitivity to several drugs. Many of these mode of action-specific genes were not previously identified as vulnerability factors using Tn-seq approaches since they constitute essential genes and were thus not part of the transposon mutant library, highlighting the benefit of using CRISPRi for chemical genetics. We further extended the chemical genetic screens to nine strains including clinically important strains and commonly used model strains, showing both strain-specific and conserved signatures. We were able to identify conserved genes between at least 7 strains in each antibiotic condition (Figure 4C) which provides a compelling foundation for future experiments. As an example, we show that downregulation of *mutS2* and *spv_1295* renders pneumococci more vulnerable to macrolides such as azithromycin, as well as linezolid. Targeting such genes could allow us to resensitize macrolide-resistant *S. pneumoniae*, which is currently categorized as a medium level threat by the WHO^4^. A systematic combinatorial screen identified nitrofurantoin and doxycycline as synergizing with azithromycin^68^, which would be interesting to explore next.

Identifying the LiaFSR system as a hit in levofloxacin treatment in two-thirds of the strains (Figure 4C) is an intriguing result as it demonstrates that targeting this system, for instance with drugs like bacitracin that activate LiaR, can be extended to multiple strains, resulting in a clinically relevant global sensitizing target for the fluoroquinolones^37^. Such findings are a major goal of genome-wide chemogenomic studies. It also further proves the power of this approach in reliably identifying important targets across the pangenome of the pneumococcus.

While we focused on hits that were conserved in at least seven strains (Figure 4C), future work can explore strain-specific targets, in the context of clinical source, pathogenicity or resistance profiles. For example, a study using CRISPRi to compare rifampicin sensitive and resistant *M. tuberculosis* strains revealed multiple genes that were involved in driving the fitness of resistant strains^69^. Similarly, in future work we could compare the antibiotic sensitive and resistant pneumococcal strains in this panel to find genetic factors driving resistance.

The multi-strain approach addresses the shortcomings of using a single reference strain, such as D39V, to extrapolate the antibiotic response of the entire species. Drug–drug interactions are often species-and strain-specific, as demonstrated in a study that found up to 20% are strain specific^70^. One major limitation in antimicrobial compound screens is strain-specific gene variation, which confounds target selection and leads to compounds ineffective across multiple strains^71^. This highlights the need for large-scale studies to map these complexities and inform more effective treatment strategies. Indeed, using Tn-seq in nine strains of *Mycobacterium tuberculosis* also identified strain-specific signatures^72^. Also for *S. pneumoniae*, a correlation was described between overall gene essentiality, and strain background in a set of 36 different strains^18^. Additional Tn-seq studies coupled to RNA transcriptome profiling using RNA-seq, failed to identify a correlation between differential gene expression and the phenotypic importance^21^. A study in *Staphylococcus aureus* used Tn-seq to identify genes important in daptomycin stress across five strains and were able to identify some core genes shared across the strains, but also identified several strain-specific genes^73^. It will be interesting to see whether the here observed conserved pangenome-specific antibiotic stress signatures are also present across strains of other bacterial species. Recent work from our lab and others also showed mode of action specific antibiotic stress signatures using CRISPRi in *S. aureus*, suggesting that this might be the case^31,74^.

Together, our findings reveal genome-wide genetic factors that modulate antibiotic sensitivity, highlighting conserved and strain-specific signatures that can be explored. This distinction is critical for identifying universal drug targets as well as strain-specific therapeutic strategies, facilitating the development of both broad-spectrum and precision treatments.

## Materials and Methods

### Bacterial strains, culture conditions, and antibiotic stocks

The strains studied in this study are listed in Table 1 and Supplementary Data 15. All *S. pneumoniae* experiments were performed in plain C+Y medium at 37°C, unless otherwise noted^75^. C+Y medium was supplemented with choline chloride (10 mg/ml) when necessary. For induction of dCas9, anhydrotetracycline (100 ng/ml), IPTG (40 µM), or doxycycline (20 ng/ml) was used appropriate. Antibiotic selection for *S. pneumoniae* was performed on spectinomycin (100 μg/ml), tetracycline (0.5 μg/ml), gentamicin (40 μg/ml), or trimethoprim (10 μg/ml). *S. pneumoniae*-80°C freezer stocks for were prepared by culturing strains to an OD_595_ = 0.4. The culture was centrifuged (14000 × *g*, 10 minutes) and the pellet resuspended in fresh medium supplemented with 15% glycerol.

*E. coli* cultures strains were grown in lysogeny broth (LB) at 37°C with agitation. LB was supplemented with spectinomycin 100 μg/ml when appropriate.

All antibiotics were acquired from Sigma-Aldrich.

### Genomic analyses

Genomic sequences (Table 1) from the studied strains were annotated *de novo* as described before, using a customized version of Prokka (v1.14.6)^28,76^. For this, the “-c” flag for Prodigal (v2.6.3)^77^ was removed, enabling annotation of open reading frames at the start or end of a contig without a matching start-or stop codon. Further annotation was enabled by using ARAGORN (v1.2.38)^78^ for transfer RNAs, Infernal (v1.1.4)^79^ and Rfam^80^ for non-coding RNAs, MinCED (v0.4.2)^81^ for the identification of CRISPR arrays, and HMMER3 (v3.3.2)^82^ for protein similarity search. The previous high-detail annotation of D39V was used as reference during the annotation process^29^, which has since been further refined^28^. Prokka-annotated genomes were analyzed for matching genomic content using Panaroo (v1.2.10)^83^ using --clean-mode “strict”, and –remove-invalid-genes. COG clusters were assigned based on the D39V genome (CP027540.1) using EggNOG^84^.

### sgRNA library construction

For the design of the sgRNA libraries, a previously described pipeline was used, in conjunction with the Prokka-based annotation^48^. The designed sgRNA sequences were extended by adding BsmBI (Esp3I isoschizomer) restriction sites, strains-specific primer sites, and universal primer sites to each side, resulting in a sequence of 142 nucleotides per sgRNA. The primer sites were based on 25-nucleotide long orthogonal sequences designed elsewhere^85^. The designed oligonucleotides were synthesized as a pooled library of single stranded DNA molecules by Twist Bioscience (South San Francisco, California, United States of America). The pooled oligonucleotides were resuspended in Buffer EB (10 mM Tris-HCl, pH 8.0; Qiagen) and stored at-20°C until further use. Strain-specific portions of the pooled library were amplified for use with strain-specific primer combinations (OVL6482-OVL6487, OVL6496-OVL6499; OVL9383-OVL0384; OVL9632-OVL9633 (Supplementary Data 16)), after which the 92 base pair fragments were purified using the Monarch PCR & DNA Cleanup kit (5 μg; T1030), using the oligonucleotide protocol. For each strain of interest, a strain-specific mCherry-encoding pPEPZ/pPEPY plasmid was constructed using Golden Gate cloning, containing native ZIP/CIL-locus flanking sequences^86^. For all strains the ZIP locus was selected, except for the ZIP-locus deficient SN75752, for which the CIL-locus was selected as integration site (Supplementary Data 15, Supplementary Data 16, Supplementary Data 17). These plasmids were linearized using EcoRV-HF, after which the backbone was amplified (OVL6185/OVL6186). Purified backbone and sgRNA oligonucleotides were used in a ratio-optimized Golden Gate reaction in a high throughput cloning step, utilizing Esp3I and T4 ligase. Resulting reactions were dialyzed and transformed into freshly grown *Escherichia coli* Stbl3 cells that were extensively washed in deionized water, using electroporation (2500 V), after which the cells were recovered Super Optimal broth with Catabolic repressor, and plated on LB agar supplemented with spectinomycin. After overnight growth at 37°C, colonies were enumerated, to ensure a low number of mCherry-positive colonies and a minimal coverage of 20 times the designed strain-specific sgRNA library, after which the cells were scraped from the plates, and stored at-80°C in LB supplemented with 20% glycerol.

Subsequent miniprep-obtained plasmids were transformed into strains constructed to harbor a tetracycline-like molecule inducible dCas9 construct integrated downstream of the gene encoding β-galactosidase BgaA. This construct included a constitutively expressed *tetM* tetracycline resistance gene, to confer resistance to the inducer, if the strain was not already resistant to tetracycline-like antibiotics. For transformation, a standard pneumococcal transformation protocol utilizing strain-appropriate synthetic competence-stimulating peptide (CSP) was used^87^. After selective growth on Colombia agar supplemented with 3% defibrinated sheep blood and spectinomycin (100 μg/ml), colonies were enumerated to ensure at least a 20 times coverage of the designed library, after which colonies were scraped from the plates using C+Y medium supplemented with spectinomycin (100 μg/ml) and 20% glycerol, and stored at-80°C.

For the cloning of sgRNAs, two partially reverse complementary oligonucleotide primers were ordered to encode the sgRNA sequence and the Esp3I-produced four-nucleotide overhangs (OVL6808-OVL6909, OVL10371-OVL10372, OVL12258-OVL11265). After annealing in TEN buffer (10 mM Tris-HCl, 1 mM EDTA, 0.1 M NaCl, pH=8.00)^88^, sgRNAs were assembled into the prepared linear pPEPZ/pPEPY fragments using a Esp3I-mediated Golden Gate reaction, transformed into *E. coli* Stbl3, after which the resulting plasmids were transformed into their specific dCas9-carrying pneumococcal strains.

### Chemical genetics using the D39V operon-level CRISPRi-seq library

To determine sub-inhibitory antibiotic concentrations to use in the chemical genetic screening using the D39V operon-level library, cells were first inoculated from a-80°C freezer stock 1:100-fold and grown to an OD_595_ = 0.1. This culture was then further diluted 100-fold into medium containing a range of concentrations spanning the approximate MIC of the relevant antibiotic, into a 96-well flat-bottom plate. This plate was incubated at 37°C, and the OD_595_ was measured every 10 minutes using the Tecan Infinite 200 plate reader. Growth curves were plotted using BactExtract^89^. For every antibiotic, a sub-inhibitory concentration that resulted in a distinct growth defect compared to a control strains were determined from these growth plots (Supplementary Data 1).

To perform the chemical genetic screen, the pooled D39V operon-level was diluted 100-fold in C+Y medium in separate biological replicates, and grown to an OD_595_ = 0.1^32,33^. Fresh cultures were then inoculated 1:10, IPTG added when appropriate (i.e. the inducer to activate the dCas9 system), and grown to an OD_595_ = 0.1. The IPTG-induced and non-induced cultures were each diluted 150-fold into 15 ml fresh medium in a 15 ml Falcon tube, under one of four conditions: I) minus IPTG, minus antibiotic; II) plus IPTG, minus antibiotic; III) minus IPTG, plus antibiotic; or IV) plus IPTG, plus antibiotic. Conditions I and III will yield the baseline composition of the library without expression of dCas9. In condition II and IV the dCas9 system is expressed, causing depletion of the cells of the library in which the sgRNA targets dCas9 to genetic elements that are important for survival. Cultures were grown to OD_595_ = 0.4 (∼11 generations), and centrifuged (4000 × *g*, 10 minutes, 4°C). Cells were washed once in phosphate buffered saline (pH = 7.4) and stored at-80°C for genomic DNA (gDNA) extraction.

For gDNA extraction, the frozen cell pellet was resuspended in 50 µl of water and half of the volume taken for gDNA extraction. The aliquot was resuspended in 800 µl of Nuclei Lysis solution (Promega) supplemented with 0.05% SDS, 0.025% deoxycholate (DOC), and 200 µg/mL RNase A. This suspension was incubated at 37°C for 20 minutes, then at 80°C for 5 minutes to lyse cells, then at 37°C for 10 minutes. After addition of Protein Precipitation Solution (250 µl; Promega), the sample was vortexed vigorously, then incubated on ice for 10 minutes. Samples were centrifuged (14000 × *g*, 10 minutes, 4°C) to pellet the precipitated proteins. The supernatant was transferred into 600 µl isopropanol to precipitate the gDNA, which was then collected by centrifugation (14000 × *g*, 10 minutes). The gDNA pellet was washed once in 70% ethanol, air-dried, and resuspended in molecular grade water. The resulting gDNA was stored at-20°C. Extracted gDNA was used for the amplification of remaining sgRNAs according to previously published protocols, which were subsequently sequenced using an Illumina MiniSeq sequencing system using a 2 x 150 high-output kit, according to a custom sequencing protocol^48^.

### Timepoint sampling of gene-level CRISPRi-seq libraries

Stored pneumococcal libraries were inoculated in 15 ml medium (supplemented with doxycycline when appropriate) 1:100 (theoretical OD of 0.003), and grown to an OD of 0.3 (∼7 generations) at 37°C. cultures were passed into fresh medium with the same conditions 1:100, and leftover cells were centrifuged (3000 × *g*, 10 minutes, 4°C), washed in PBS (pH = 7.4), and stored at-80°C. After 21, all cells were stored after centrifugation and washing. Stored cells were used for gDNA extraction according to the methods described for the D39V operon-level CRISPRi-seq library. The amplified sgRNAs were sequenced using an Illumina NovaSeq 6000 sequencing system.

### Testing the effect of choline chloride on cell autolysis

To assess the impact of choline chloride on *S. pneumoniae* autolysis, C+Y medium was supplemented with 10mg/ml of choline chloride and growth was monitored by OD_595_. It was confirmed that the addition of choline chloride inhibited autolysis and allowed for the overnight growth of cultures^90^, even in the presence of amoxicillin (Supplementary Figure 11A). To test the effect of choline chloride and overnight growth on antibiotic gene essentiality profiles, a test screen was conducted using the D39V and PBCN0272 gene-level CRISPRi libraries. The libraries were grown overnight in sub-inhibitory concentrations of amoxicillin. The resulting significant gene hits were compared to the previous amoxicillin screen in the D39V operon-level library and found to be similar (Supplementary Figure 11B, Supplementary Data 18). This suggested this modified approach could be utilized, allowing for these screens to be performed in a high-throughput fashion.

### Chemical genetic screens using gene-level CRISPRi-seq libraries

To determine sub-inhibitory antibiotic concentrations to use in the screening, each pneumococcal library was inoculated and grown to an OD_595_ = 0.1. Cultures were diluted 150-fold into fresh medium 10mg/ml choline chloride and a range of antibiotic concentrations. The OD_595_ was monitored over 20 hours. Sub-inhibitory antibiotic concentrations were determined as described for the D39V operon-level CRISPRi-seq library (Supplementary Data 10).

To perform the chemical genetic screen, biological replicates of 4 ml were inoculated 1:20 with the CRISPRi-seq library of each respective strain and grown until OD_595_ = 0.15. Then, 4.5 ml fresh medium containing choline chloride was inoculated 1:30 with the grown library cells. Cultures were supplemented to one of four conditions: I) minus aTc, minus antibiotic; ii) plus aTc, minus antibiotic; III) minus aTc, plus antibiotic; or IV) plus aTc, plus antibiotic. Cultures were grown for 18 hours, after which cultures were centrifuged (3000 × *g*, 20 minutes, 4°C) to pellet cells, supernatant removed and stored at-80°C for gDNA extraction.

From the stored pellets, gDNA was isolated using the FastPure Bacteria DNA isolation mini kit (DC103-01, Vazyme), according to the manufacturer’s protocol, with the following minor alterations. Frozen bacterial cell pellets were resuspended in buffer GA supplemented to include 0.05 % SDS, 0.025 % DOC, and 200 µg/ml RNase A. After incubation at 30°C for 10 minutes, proteinase K was added, and the sample was further incubated at 37°C for 10 minutes. After addition of buffer GB, the mixture was further incubated for 10 minutes at 80°C, after which the protocols standard column-based workflow was followed. Genomic DNA was eluted in 50 µl of molecular grade water. DNA was stored at-20°C Extracted gDNA was used for the amplification of remaining sgRNAs according to previously published protocols, which were subsequently sequenced using a AVITI sequencing system.

### Strain construction

To construct strains harboring the tetracycline-like molecule inducible dCas9 construct, we amplified approximately 1000 bp portions up-and downstream of the targeted integration site. Up-and down-stream portions were fused to the dCas9-carrying construct using Golden Gate reactions. This construct included a constitutively expressed *tetM* tetracycline resistance gene, to confer resistance to the inducer, if the strain was not already resistant to tetracycline-like antibiotics. After, the product was transformed into the wild-type strains through standard pneumococcal transformation protocols using strain-appropriate CSP^87^. In contrast to the D39V operon-level library, all gene-level libraries were constructed to have the dCas9-carrying construct integrated downstream of *bgaA*, instead of disrupting it. The oligonucleotide primers and amplified fragments for the construction of these strains can be found in Supplementary Data 16 and Supplementary Data 17.

### Construction of gene deletion mutants

Deletion mutants were generated by allelic exchange using Golden Gate assembly to fuse upstream and downstream homologous regions to an antibiotic resistance cassette. A full summary of amplified fragments and reactions can be found in Supplementary Table 17 Specifically, for deletion of the *SPV_1295* homologue in Taiwan19F, strain VL8169 (Taiwan19F + *EMCPDKAE_00810*::*aac(3)I*) was constructed. The upstream homology arm (ISU) was PCR-amplified from Taiwan19F wild type (VL124) using primers OVL12432 and OVL12474, and the downstream homology arm (ISD) was amplified from the same template using primers OVL12218 and OVL12471. The gentamicin resistance cassette (*aac(3)I)* was amplified from plasmid pVL7228 (pPEPZ-aac(3)I-mCherry-SpT) using primers OVL12473 and OVL12472. The three fragments were assembled by Golden Gate cloning using BsaI.

For deletion of the *SPV_1295* homologue in PBCN0272, strain VL8185 (PBCN0272 + *CLMNKJOF_00025*::*dfrA21*) was generated. ISU and ISD fragments were amplified from PBCN0272 wild type (VL4215) using primer pairs OVL12432/OVL12214 and OVL12217/OVL12218, respectively. The trimethoprim resistance cassette (*dfrA21)* was amplified from strain VL7928 (D39V + *SPV_1295*::*dfrA21*) using primers OVL12215 and OVL12216. The three fragments were assembled by Golden Gate cloning using Esp3I.

For deletion of *mutS2* in PBCN0272, strain VL8227 (PBCN0272 + *mutS2*::*dfrA21*) was constructed. ISU and ISD fragments were amplified from PBCN0272 wild type (VL4215) using primer pairs OVL12219/OVL12220 and OVL12421/OVL12223, respectively. The trimethoprim resistance cassette was amplified from strain VL7930 (D39V + *mutS2*::*dfrA21*) using primers OVL12222 and OVL12221. The three fragments were assembled by Golden Gate cloning using Esp3I.

### Phylogenetic analysis

A core genome single-nucleotide polymorphism (SNP) phylogeny was built according to previously published methods^28^. In short, the genomic sequences of the strains used in this study, and those from a reference dataset published previously^47^, were aligned using snippy (available from https://github.com/tseemann/snippy). Duplicate genomes strains were filtered out of the genome set from Antic *et al.* Alignments were corrected for recombination using Gubbins (v3.3.1)^91^, and the core genome SNP phylogeny was constructed using FastTree (v2.1.10)^92^. The produced phylogenetic tree was midpoint rooted using the phangorn package in R (v4.3.0)^93^.

### Differential fitness analysis

Sequencing reads from both the Illumina and Element Biosciences sequencing systems were quantified for their sgRNA content using 2FAST2Q (v2.7.8) using the following settings: minimum Phred quality score 25, number of mismatches 0, and start position of 0 or 54 (depending on the used sequencing platform)^94^. Further quality control measures to limit the effects of random noise were taken by filtering out the sgRNAs that had less than 20 reads on average in uninduced samples. Differential abundance analysis was then performed using DESeq2^95^. The interaction effect of antibiotics was assessed by comparing the differential fitness between antibiotic-exposed dCas9-induced and uninduced samples on one side, and the differential fitness between non-exposed dCas9-induced and uninduced control samples (of the same day) on the other. Log_2_FCs were shrunk using the “Adaptive Shrinkage” (ashr) package (available from github: https://github.com/stephens999/ashr).

Throughout the analyses, significant changes between compared groups were defined as an absolute log_2_FC larger than 1, and an adjusted p-value smaller than 0.05. Differences between groups were assessed using a PERMANOVA test, using the vegan R package (https://github.com/vegandevs/vegan). Post hoc testing was performed using pairwise.adonis2 (available from github: (https://github.com/pmartinezarbizu/pairwiseAdonis)). Heatmaps were drawn using ComplexHeatmap^96^. KEGG enrichment analyses were performed using clusterProfiler^97^. Upset plots were produced using UpSetR^98^.

## Data availability

The sequencing data describing the sgRNA content of the *S. pneumoniae* D39V operon level CRISPRi-library after exposure to fluoroquinolones are available in the NCBI Sequence Read Archive under BioProject accession number PRJNA1185710 from a previous publication^37^. All other sequencing data are available in the NCBI Sequence Read Archive under BioProject accession number PRJNA1347041. In addition, gene essentiality results and sgRNA sequences have been integrated for all strains (except PBCN0272 and PBCN0364) in PneumoBrowse 2^28^. The gene essentiality data (timepoint experiments, and antibiotic-exposed) are available in the information accompanying each coding feature, whilst sgRNAs can be viewed in the “Gene level” CRISPRi-track for each genome.

## Supporting information

Description of Supplementary Tables

## Acknowledgements

We thank the Lausanne Genome Technologies Facility at the University of Lausanne for their expertise using the Illumina NovaSeq 6000 and Element Biosciences AVITI sequencing systems. Strains SN75752, Spn1439-106, and Taiwan19F were a kind gift of Mark van der Linden at the University Hospital RWTH Aachen in Aachen, Germany. Strain EF3030 was a kind gift of Sven Hammerschmidt from the University of Greifswald in Greifswald, Germany. Strain BHN418 was a kind gift of Birgitta Henriques-Normark from the Karolinska Institute in Stockholm, Sweden. Strains PBCN0272 and PBCN0364 were kindly gifted by Marien de Jonge from the Radboud University Medical Center in Nijmegen, the Netherlands.

## Author Contributions

Experimental work was performed by BS, ABJ, LSM, BR, MRG, AJHC. Data analysis was performed by BS, ABJ, VdB. Manuscript writing and editing by BS, ABJ, JWV with feedback from all authors.

