## Supplementary material for "A functional genetic landscape of antibiotic sensitivity across the pneumococcal pangenome reveals conserved and lineage-specific vulnerabilities": Description of Supplementary Tables

**Description of Additional Supplementary Files**

### **Supplementary Data 1. Antibiotics used in the chemical genetic CRISPRi-seq screens tested against the D39V operon-level sgRNA library.** For each antibiotic, the class, mode of action, abbreviation, and sub-inhibitory concentration used in this study, are noted.

### **Supplementary Data 2. Raw and normalized sgRNA counts for the D39V operon-level CRISPRi-seq library screened against 15 antibiotics.** Raw counts were extracted from the sequencing reads using 2FAST2Q. Raw counts were filtered for sgRNAs that had less than 20 reads on average in uninduced samples and normalized using DESeq2.

### **Supplementary Data 3.** **D39V operon-level CRISPRi-seq library gene essentiality in the presence of antibiotics.** Results from the essentiality analyses for the D39V operon-level CRISPRi-seq library, per (antibiotic) condition. For these analyses, the IPTG-induced samples were compared to the non-induced samples within a condition. The control samples from each experimental day were treated separately. The resulting log_2_FC values were shrunk using ashr.

### **Supplementary Data 4.** **D39V operon-level CRISPRi-seq library interaction data in the presence of antibiotics.** Interaction analyses for the D39V operon-level CRISPRi-seq library per antibiotic. For these analyses, the fitness values of the antibiotic-exposed samples were compared to the C+Y control samples of the same experimental day.

### **Supplementary Data 5. Designed gene-level sgRNAs for the nine phylogenetically diverse *S. pneumoniae* strains.** Using a recently published pipeline^40^, we designed sgRNAs targeting every predicted gene in the genomic sequences of the strains of interest. For each strain, the designed sgRNAs and their properties are indicated.

### **Supplementary Data 6. Raw counts for the nine gene-level CRISPRi-seq libraries screened at different timepoints (7, 14, 21 generations).** Raw counts were extracted from the sequencing reads using 2FAST2Q. The underlying sequencing reads can be acquired from the NCBI under BioProject accession codes PRJNA1185710 and PRJNA1347041.

### **Supplementary Data 7. Normalized counts for the nine gene-level *S. pneumoniae* CRISPRi-seq libraries screened at different timepoints (7, 14, 21 generations).** Raw counts were filtered for sgRNAs that had less than 20 reads on average in uninduced samples and normalized using DESeq2. The underlying sequencing reads can be acquired from the NCBI under BioProject accession codes PRJNA1185710 and PRJNA1347041.

### **Supplementary Data 8.** **Gene-level CRISPRi-seq libraries essentiality results screened at different timepoints (7, 14, 21 generations).** Results from essentiality analyses for the nine gene-level CRISPRi-seq libraries, per timepoint (7, 14, or 21 generations). For these analyses, the doxycycline-induced samples were compared to the non-induced samples within a condition. The log_2_FC values were shrunk using ashr.

### **Supplementary Data 9.** **Qualitative comparison in gene essentiality of protein-encoding genes between previously performed Tn-seq studies, and the work presented here.** For D39V, Taiwan19F, and TIGR4, previously performed studies determined gene essentiality using Tn-seq. The results from the 21 generations timepoint in this study were compared to the previously published results in a qualitative approach, without consideration of generation time or medium. If more than two gene essentiality classifications were present in the Tn-seq studies, only the strictest classification was considered for the qualitative comparison. Locus tags were matched between studies using Panaroo.

### **Supplementary Data 10. Sub-inhibitory concentrations of the antibiotics used in the multi-strain chemical genetic CRISPRi-seq screens.** For each antibiotic, the sub-inhibitory concentration (in μg/ml) per strain, as used in the chemical genetic screens is noted. These concentrations were determined in C+Y supplemented with 10 mg/ml choline chloride.

### **Supplementary Data 11.** **Interaction analyses for nine gene-level libraries in presence of four different antibiotics.** Interaction analyses for the nine gene-level CRISPRi-seq library per antibiotic. For these analyses, the fitness values of the antibiotic-exposed samples were compared to the C+Y control samples.

### **Supplementary Data 12.** **Raw counts for the nine gene-level CRISPRi-seq libraries cultured in the presence of four different antibiotics.** Raw counts were extracted from the sequencing reads using 2FAST2Q. The underlying sequencing reads can be acquired from the NCBI under BioProject accession codes PRJNA1185710 and PRJNA1347041.

### **Supplementary Data 13.** **Normalized counts for the nine gene-level CRISPRi-seq libraries cultured in the presence of four different antibiotics.** Raw counts were filtered for sgRNAs that had less than 20 reads on average in uninduced samples and normalized using DESeq2. The underlying sequencing reads can be acquired from the NCBI under BioProject accession codes PRJNA1185710^34^ and PRJNA1347041.

### **Supplementary Data 14.** **Essentiality analyses for gene-level CRISPRi-seq libraries in presence of different antibiotics.** Essentiality analysis for the nine gene-level CRISPRi-seq libraries per condition. For these analyses, the aTc-induced samples were compared to the non-induced samples within a condition. The log_2_FC values were shrunk using ashr.

### **Supplementary Data 15.** **Genetically modified strains used in this study.** For each genetically modified strain used in this study, the in-house number, the genetic background (Table 1), description, and genotype is given. If applicable, a reference is noted.

### **Supplementary Data 16. Oligonucleotides used in this study.** For each oligonucleotide, the in-house number, the sequence (5’-3’), and a reference (when applicable) are given.

### **Supplementary Data 17.** **Overview of amplified fragments for strain construction.** The genetic modification of the strains noted in Table 1, resulted in the strains in Supplementary Data 15, the following PCRs on specified template strains were performed using specified oligonucleotide primers (Supplementary Data 16) and restriction enzymes (for Golden Gate cloning).

### **Supplementary Data 18.** **Supporting information for choline chloride analysis in D39V and PBCN0272.** Raw counts, normalized counts, essentiality calls, and interaction calls for the gene-level libraries D39V and PBCN0272 CRISPRi libraries cultured in C+Y supplemented with amoxicillin, with and without choline chloride. Raw counts were extracted from the sequencing reads using 2FAST2Q. Raw counts were filtered for sgRNAs that had less than 20 reads on average in uninduced samples and normalized using DESeq2. The log_2_FC values were shrunk using ashr. The underlying sequencing reads can be acquired from the NCBI under BioProject accession codes PRJNA1185710^34^ and PRJNA1347041.
