## Supplementary Figures for "A functional genetic landscape of antibiotic sensitivity across the pneumococcal pangenome reveals conserved and lineage-specific vulnerabilities"

**Supplementary Information**

**Supplementary Figure 1.** **Interaction scatter plots for D39V operon-level library exposed to 15 different antibiotics.** The D39V operon-level library was exposed to 15 different antibiotics. For each analysis, ashr-shrunk and scaled log_2_FCs, and adjusted P-values are plotted. Data points with an absolute interaction log_2_FC > 1, and an adjusted-P-value < 0.05 were considered to have a significant different fitness between control and antibiotic-exposed samples and are colored in orange. Labeling of datapoints is arbitrary. Full results can be found in Supplementary Table 4.

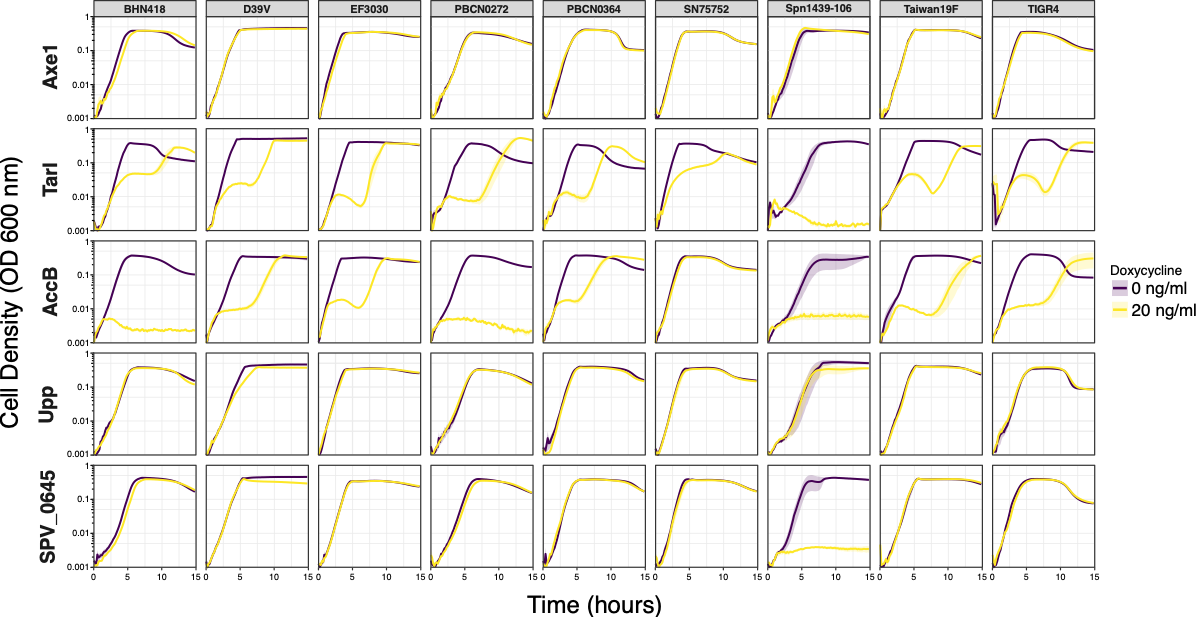

**Supplementary Figure 2. Strain-specific gene essentiality in phylogenetically diverse *S. pneumoniae* strains.** The functionality of the CRISPRi system and induction of dCas9 was confirmed by cloning of an sgRNA targeting the essential *tarI* gene. To control for non-specific effects and effects by the inducer (doxycycline), a strain carrying a control sgRNA targeting a non-essential gene (*axe1*). Strain-specific gene-essentiality was confirmed through cloning of sgRNAs targeting *accB*, *upp*, and (the ortholog of) SPV_0645.

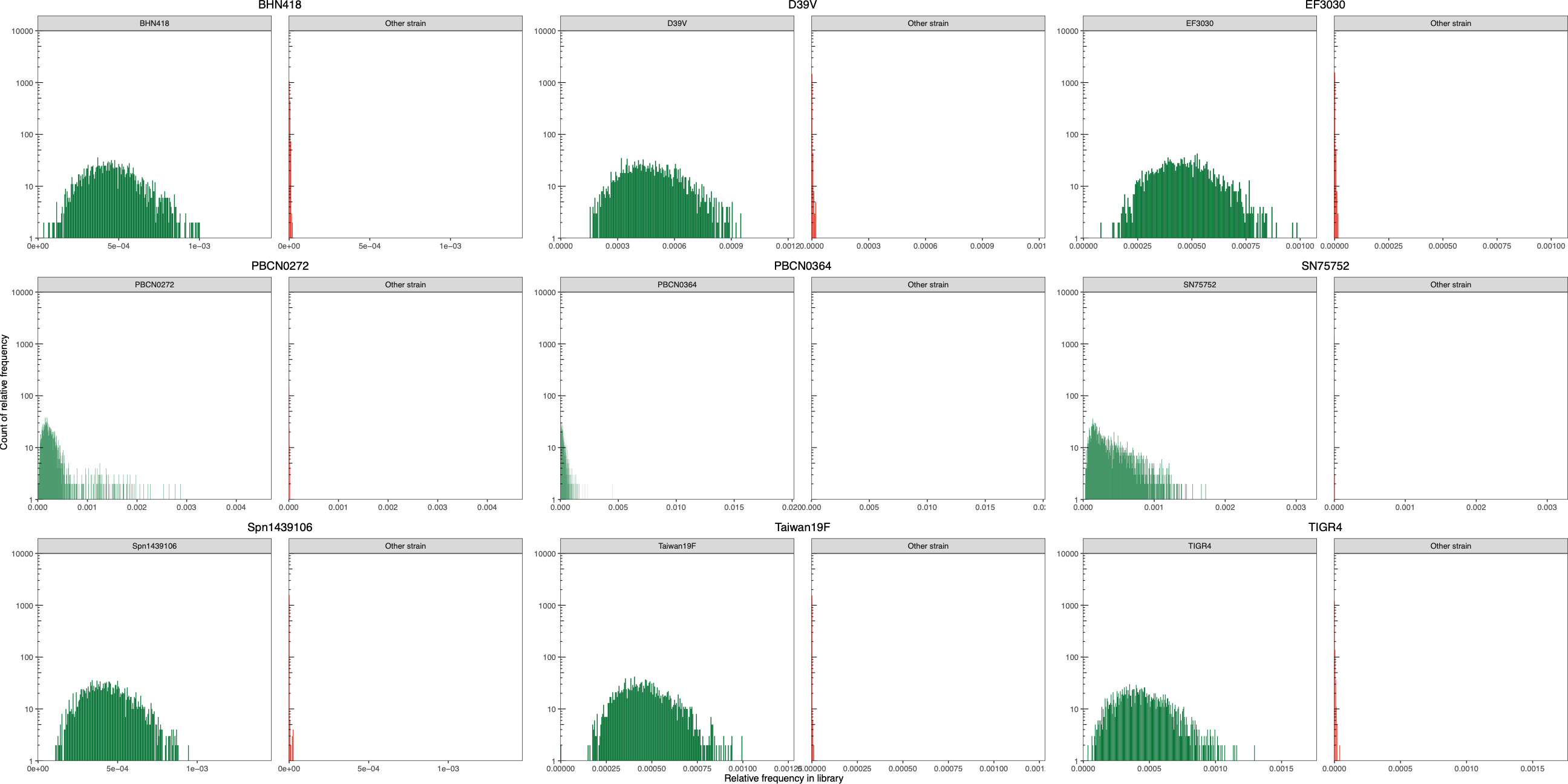

**Supplementary Figure 3. Distribution of sgRNA frequencies in cloned CRISPRi-seq libraries.** The distribution of strain-specific and strain-non-specific sgRNAs shows the specificity of the developed Golden gate-based cloning strategy.

**Supplementary Figure 4. Differential fitness in gene-level *S. pneumoniae* CRISPRi-seq libraries.** For each library, the differential fitness of each sgRNA (gene) was quantified through differential abundance analyses at three timepoints. For each sgRNA, the ashr-shrunk log_2_FCs, and the adjusted P-values are plotted. Data points with an absolute log_2_FC > 1, and an adjusted-P-value < 0.05 were considered to have a significant different fitness in doxycycline-induced samples compared to uninduced samples and are colored in orange. Labeling of datapoints is arbitrary. Full results can be found in Supplementary Table 8.

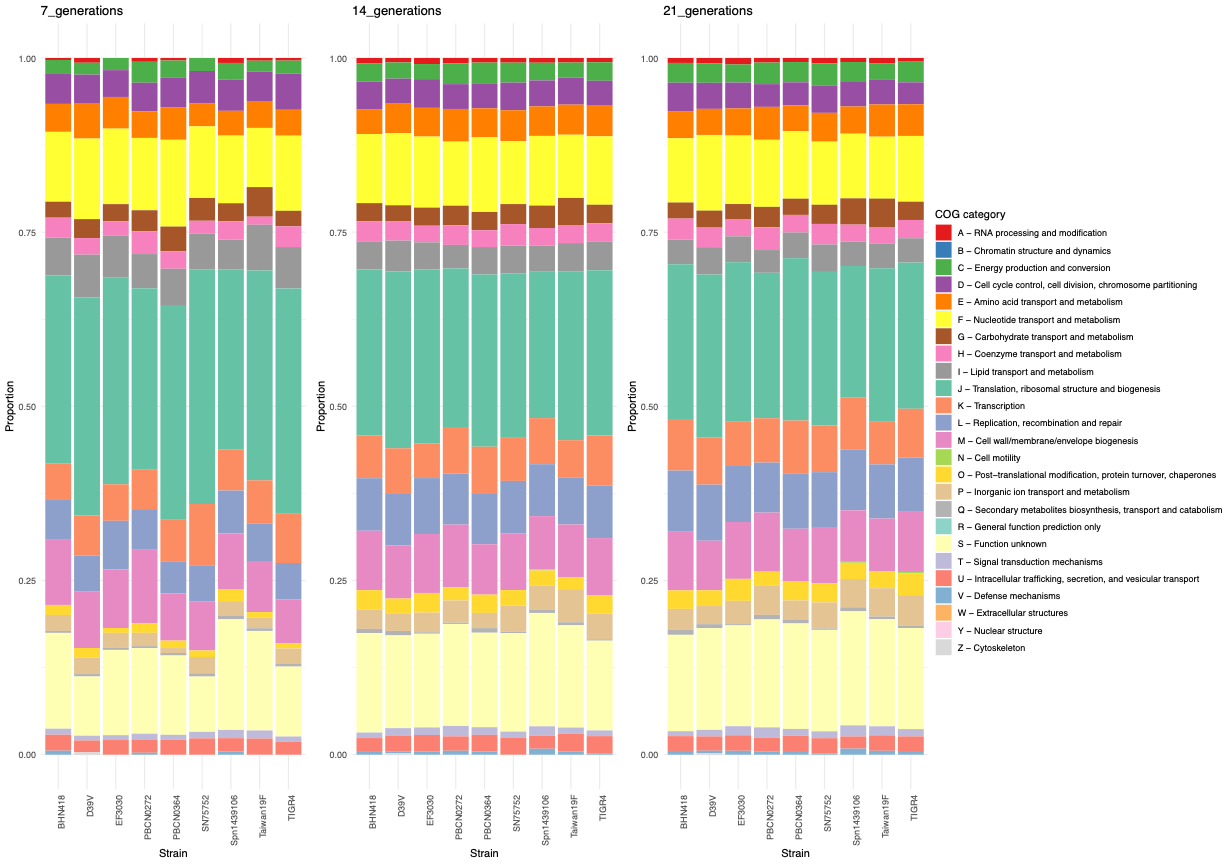

**Supplementary Figure 5. COG category analysis for the core genes determined to have a significant differential fitness difference at each timepoint.** For each library, the COG groups for the genes in the core genomes were determined at each timepoint, based on the manual annotation of the D39V genome.

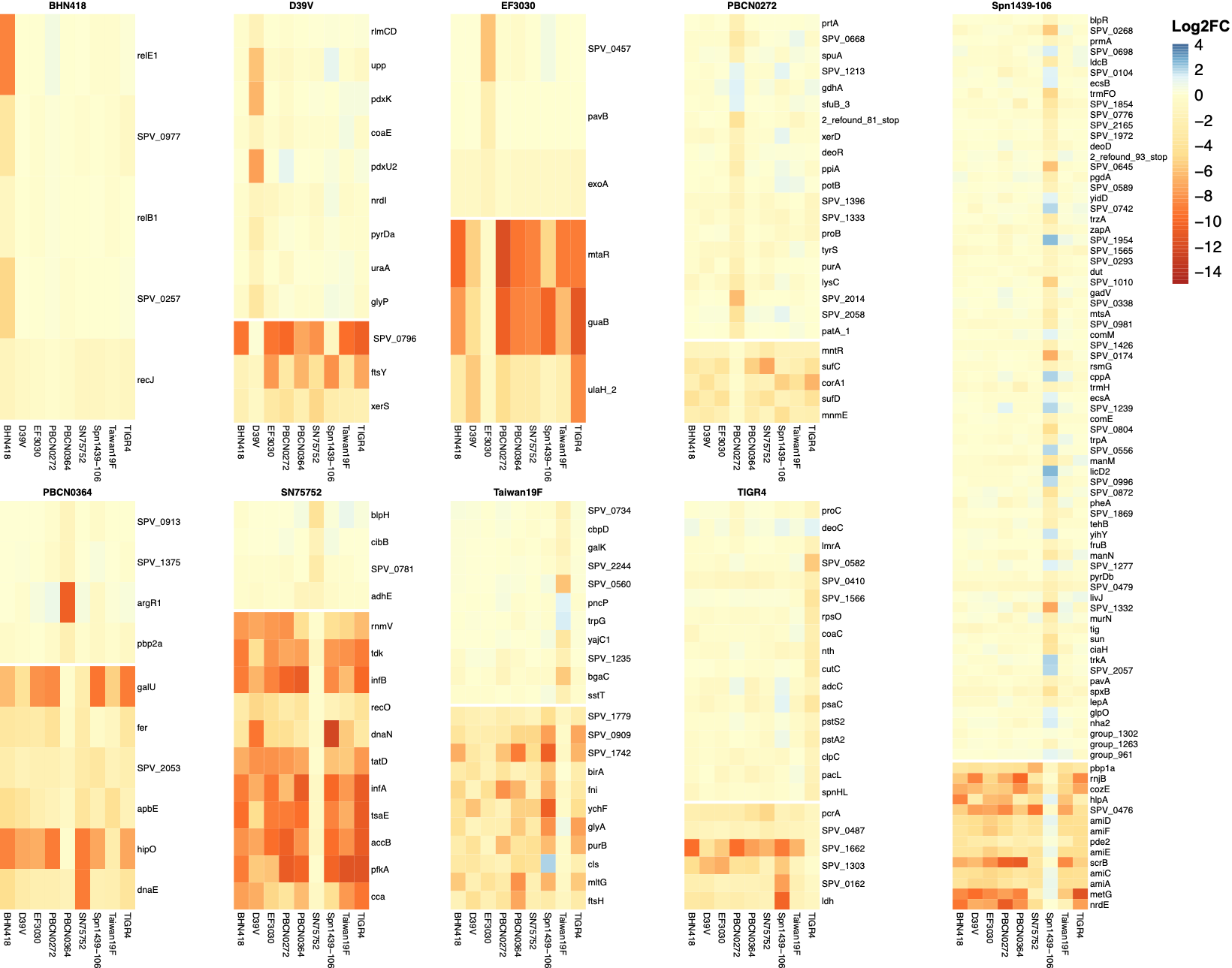

**Supplementary Figure 6. Strain-specific differential fitness after 21 generations of doxycycline-induced growth.** For each strain, the genes that show unique (non-)significant differential fitness to that strain are displayed. Gene annotations are the result of Panaroo analysis, with priority for the D39V reference annotation^27^.

**Supplementary Figure 7. Gene essentiality in phylogenetically diverse strains exposed to antibiotics.** For each combination of antibiotic and strain in the multistrain analysis, the gene essentiality after overnight growth in C+Y supplemented with choline chloride is given. For each library, the differential fitness of each sgRNA (gene) was quantified through differential abundance analyses. For each sgRNA, the ashr-shrunk log_2_FCs, and the adjusted P-values are plotted. Data points with an absolute log_2_FC > 1, and an adjusted-P-value < 0.05 were considered to have a significant different fitness in doxycycline-induced samples compared to uninduced samples and are colored in orange. Labeling of datapoints is arbitrary. Full results can be found in Supplementary Table 14.

**Supplementary Figure 8. Interaction scatter plots for the multistrain gene-level libraries exposed to four different antibiotics.** The gene-level libraries were exposed to four different antibiotics. For each analysis, ashr-shrunk and scaled log_2_FCs, and adjusted P-values are plotted. Data points with an absolute interaction log_2_FC > 1, and an adjusted-P-value < 0.05 were considered to have a significant different fitness between control and antibiotic-exposed samples, and are colored in orange. Labeling of datapoints is arbitrary. Full results can be found in Supplementary Table 11.

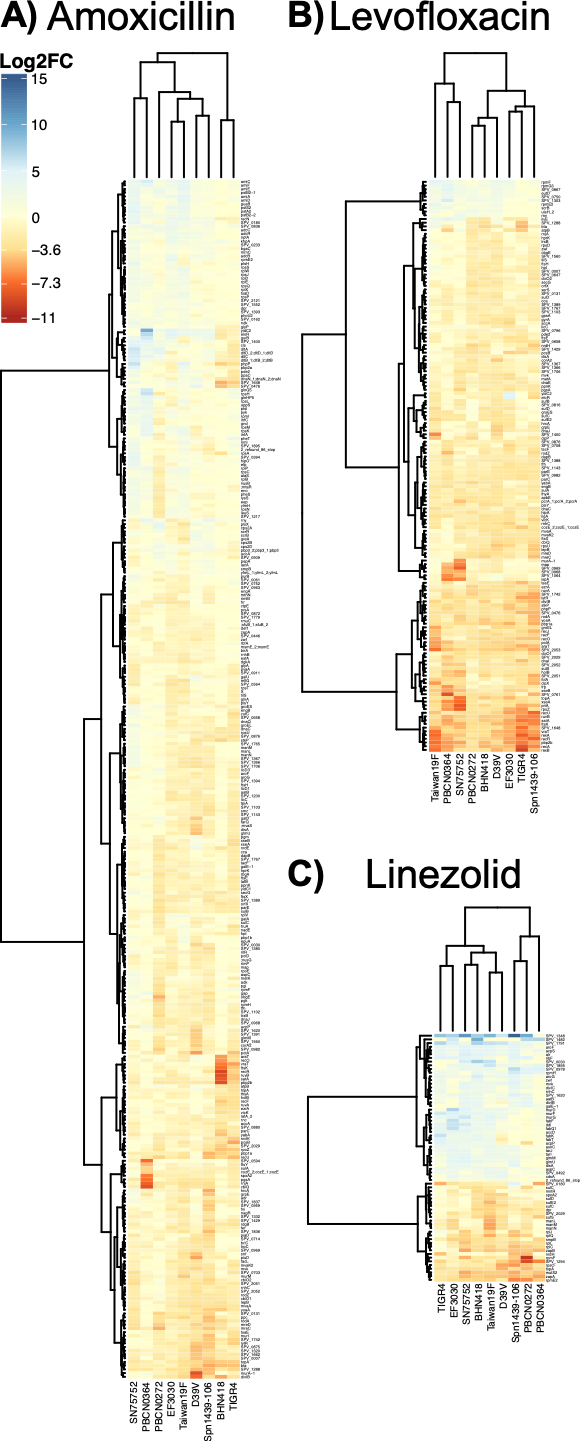
**Supplementary Figure 9. Differential gene fitness under influence of A) amoxicillin, B) levofloxacin, and C) linezolid.** For each antibiotic, the genes that displayed differential gene fitness in at least 3 antibiotics are given. Gene annotations are the result of Panaroo analysis, with priority for the D39V reference annotation^27^.

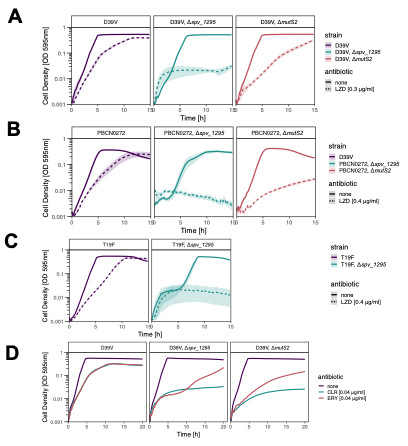

**Supplementary Figure 10. Growth of *Δspv_1295* and Δ*mutS2* mutants in the presence of linezolid and other macrolides. A)** Growth of D39V mutants treated with sub-inhibitory concentrations of linezolid (LZD). **B)** Growth of PBCN0272 mutants treated with sub-inhibitory concentrations of linezolid. **C)** Growth of T19F mutant treated with sub-inhibitory concentrations of linezolid. **D)** Growth of D39V mutants treated with sub-inhibitory concentrations of macrolides clarithromycin and erythromycin.

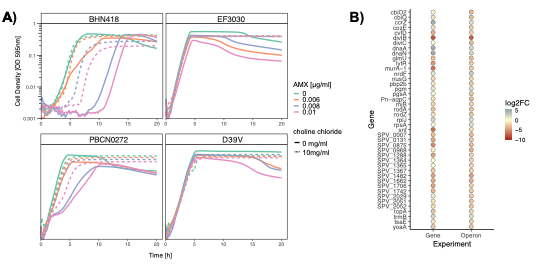

**Supplementary Figure 11. Choline chloride prevents pneumococcal autolysis enabling overnight antibiotic screening and does not affect gene essentiality profiles.** **A)** Representative growth curves of four *S. pneumoniae* strains treated with a range of amoxicillin (AMX) concentrations, in the presence and absence of 10 mg/ml of choline chloride. **B)** Comparative bubble plot between genes that had a significant fitness effect in the presence of amoxicillin between two experimental set-ups: the D39V gene-targeting library screened against amoxicillin overnight in the presence of choline chloride (experiment “Gene”) and D39V operon targeting library screened in standard C+Y (“Operon”).
